# RAPID HEMATOCRIT ESTIMATION USING A FOLD-CREASE INDUCED FAST FLOWING PAPER SENSOR

**DOI:** 10.1101/2024.01.27.577541

**Authors:** Amaan Dash, Manikuntala Mukhopadhyay, Jyoti Shaw, Maitreyee Bhattacharya, Sunando DasGupta

**Author notes:** Physical Chemistry and Soft Matter, Wageningen University and Research, Stippeneng 4, 6708 WE Wageningen, The Netherlands.

## Abstract

Increased evaporative losses and flow obstructions can present substantial impediments to current paper analytical devices (µPADs) for efficient on-site testing of biological fluids. Strategic enhancements in wicking rates of paper may thereby counter these limitations and enable on-demand healthcare monitoring. Therefore, herein we have leveraged the features of paper fold-crease regions, for the very first time, and developed a novel fast-flowing platform using laser printing to accelerate fluid flow through paper. A series of extensive experiments have been conducted to optimize the design and maximize wicking rates of µPADs for smaller liquid volumes, making it well-suited for analysing biofluids. The investigation delves into structural alterations within the creased regions, employing both static and dynamic force application strategies. A first-generation Washburn type model in excellent agreement with the experimental findings is developed, providing a comprehensive insight into the fundamental physics involved. Finally, the folded channels are utilized for a distance-based hematocrit sensor employing grade-1 filter paper at very low-cost, simplified fabrication, lesser sample volume and faster analysis. The findings of this work unveil a plethora of potentialities for employing paper and paper folds to develop affordable medical devices with advanced analytical functionalities, specifically tailored for the resource-constrained settings.

**Graphical Abstract:** 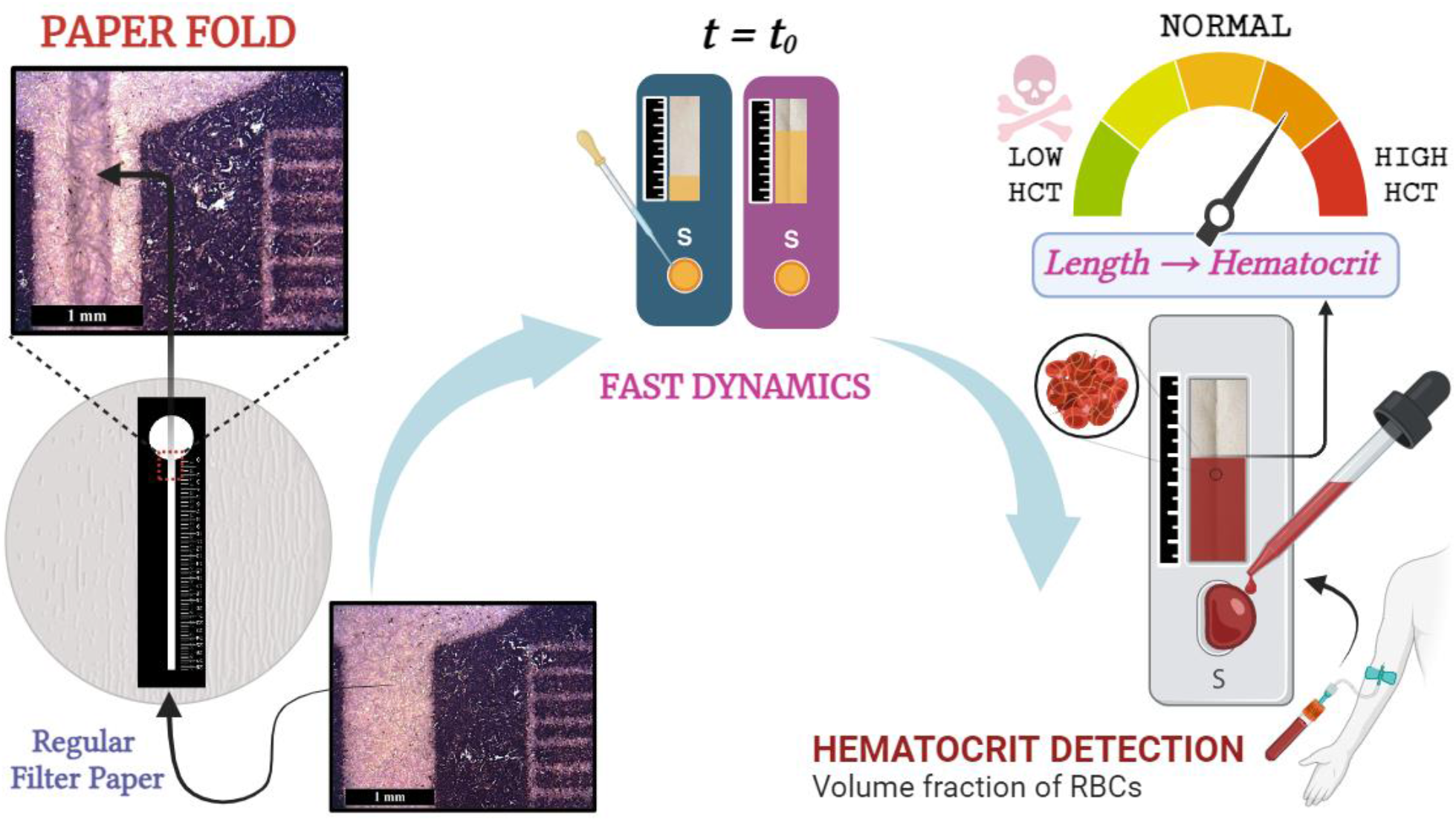

## 1. Introduction

Paper-based devices has shown immense potential for fluid manipulation in multifaceted bio-microfluidic applications [1], owing to its numerous advantages such as capillary fluid transport, minimal sample requirement, convenient disposal, biocompatibility, flexibility, and accessibility [2]. Over the last decade, it has been utilised extensively to develop point-of-care diagnostic devices for cost effective, accurate, and user-friendly testing in resource constrained settings [3]. Their material and fabrication cost have also been minimized by implementing printing techniques for large-scale production, e.g., laser [4] and wax printing [5]. However, certain limitations of paper devices (µPADs) such as fixed pore dimensions and imprecise flow control, could have an impact on the accuracy and reproducibility of certain assays, restricting their utility for on-field testing [6],[7]. Numerous techniques to control timing and sequence of fluid transport relied primarily on delaying fluid flow, by manipulating channel geometry [8], patterning hydrophobic barriers [9], control and sequential release through reservoirs and valves [10]. However, delaying flow might cause additional concerns of clogging, non-specific absorption and more critically higher evaporative loss and prolong assay time, posing difficulties for rapid and real-time monitoring of biofluids. Hence, alternative flow controlling approach, that involve accelerating fluid flow through paper, is of utmost importance to improve the accuracy, efficiency, and overall performance of the device.

In order to achieve precise flow regulation in porous media, understanding dominant factors affecting the flow is imperative. The fluid transport through paper network is governed by a complex combination of interfacial tension, capillary pressure, and other spatiotemporal parameters. The one-dimensional (1-D) wetted length for single-phase liquid transport in paper, neglecting evaporation loss and inertial effects, is given by the Lucas-Washburn equation as [11],[12]:

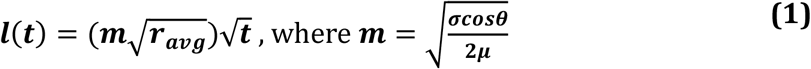

Here ***m*** is influenced by the contact angle ***θ***, surface tension ***σ*** and fluid viscosity ***µ***. Given that *θ* remains constant for a paper-fluid pair, *σ* and *μ* are properties specific to fluid, the manipulation of average pore radius ***r***_***avg***_ could be utilised to alter wicking rate through paper. Therefore, all the previous studies which aimed to accelerate the flow through paper mainly focused on bypassing a significant portion of the liquid and directing it outside the porous network in some larger external capillary. The first implementation of such fast-flowing paper channel was documented back in 2012 by incorporating flexible solid film gaps [13] and subsequently numerous techniques have been devised, including wicking through multiple layers of paper [14],[15] creating grooves [16] and hollow channels [17]. However, most of these devices requires a large sample volume, and the fabrication is rather complex and expensive. Further, making grooves and hollow channels can compromise the structural integrity of the paper, making fluid management in these devices extremely difficult. All these constraints severely limit their widespread use and adoption for the design of user-friendly and portable analytical devices.

Paper folding is also an alternate well-established technique for modulating the flow through paper-based networks. It has been applied to operate valves [13] and built origami-based designs [18]. But interestingly, the inherent paper fold line itself also has the ability to generate an enhanced capillary network within an already existing paper channel, whose capabilities still remain unexplored. These fold lines could serve as potential faster pathways for fluid manipulation, mixing and interaction for precise distribution of samples and reagents. Furthermore, it can enhance the sensitivity of the tests by enabling improved analyte access to detection zones or by promoting interactions with the substrate. In addition to all these features, it could also present several advantages in terms of fabrication, cost-effectiveness, and easy integration with patterned paper designs, therefore overcoming many of the limitations of existing fast-flowing devices. Therefore, it is imperative to investigate the structural attributes and fluid dynamics within these fold lines in order the enhance the performance of the existing µPADs.

One of the key benefits of utilizing larger pores is that it reduces the likelihood of sample retention within paper matrix, which is especially important when evaluating biological fluids having suspended cellular constituents, e.g. cerebrospinal fluid (CSF), Lymph, Synovial Fluid and Blood. Blood is a body fluid that can be characterised by metrics such as hematocrit (Hct), haemoglobin (Hgb), cell count etc [19]. The Hematocrit (Hct) is the volume fraction of red blood cells (RBCs), with normal levels in men ranging from 41-50% and 36-44% in women. Its estimate is critical for diagnosing conditions such as anaemia, polycythaemia, dehydration, and various other blood related disorders [20]. Previously a distance-based approach for estimating whole blood Hct was established using regular chromatography paper [6], but the correlation was highly inconsistent and adopting a fast-flowing paper device could potentially improve its length estimation accuracy. However, existing fast-channels have limited utility for such biofluid analysis as they cannot be implemented into devices using smaller liquid volumes and immobilised detection reagents[21]. Consequently, investigating these fold-induced fast-channels is crucial in developing efficient platforms for performing these sorts of paper based biological assays.

In this study, we leverage the features of paper fold to establish a novel fast-flowing paper platform for point-of-care hematocrit estimation using distance-based approach. This ability of paper to undergo complex folding to accelerate fluid flow retaining its structural integrity, together with their potential implications in advanced biomedical diagnostics is explored. The flow behaviour of fold introduced paper channels have been extensively investigated for the first time and the fold making technique is standardized. A predictive study, modifying the Lucas-Washburn equation has been proposed to support the flow physics and the characteristics are validated experimentally. In addition, the developed fast-flowing device is applied to improve the performance of previously established length-based hematocrit sensor with no additional cost, incorporating a simplified fabrication technique, requirement of lesser sample volume and faster analysis. The Hct results were benchmarked using an Automated Haematology Analyser and the device could successfully differentiate healthy and anaemic samples, hence facilitating low-cost point-of-care anaemia detection in resource constrained environment.

## 2. Materials and Methods

### 2.1. Fast-Flowing Channel Fabrication

The fast-flowing paper channels were fabricated using a two-step procedure. (1) The designs were patterned on Grade-1 filter paper (optimized for laser printing method) using a commercially available laser printer (HP Laser Pro 400 series) (refer **ESI. 1**.) (2) The implementation of a straight fold in the paper device involved aligning a ruler parallel to the edges of the channel, on which the fold will be implemented in a precise manner and aligned accurately. The smooth edges of the ruler effectively preclude the risk of tearing or damaging of paper fibres during the process. A predetermined and optimized force was applied along its length to introduce the fold throughout the channel. An identical procedure was repeated for each device on the filter paper. Further, the device was positioned on the hot plate (Tarson Hot top Digital MC-02) with a steel block on top (dia. 110 mm and thickness 27 mm), preheated to 180°C, for 30 sec, to ensure that the final device is straight and ready for the dispensation of the sample. The channels once prepared were stored under ambient condition until further use, without any changes in the wicking properties.

The force applied to introduce fold can be categorised into 2 broad types based on their nature: static force and dynamic force, as shown in **Fig. 1**. In static force method, two blocks of varying weights: *w*_1_and *w*_2_ (*w*_1_ > *w*_2_), were positioned on top of the introduced fold for 30 sec, whereas in the dynamic force method, the force was manually exerted on the fold using a ruler. The introduced fold was imaged by positioning the section under an upright microscope (Leica DM6000M) in reflection mode using a 5X objective lens, where the dimensions and structure of the fold, with respect to channel width **(*W*)** can be clearly visualized **(Fig. 2(a)** and **Fig. 2(b))**. Scanning Electron Microscopy (SEM), (JEOL, Model JSM-7610F) was employed to provide an estimate of the fold wavelength (***λ***) and average pore dimensions **(*r***_***f***_) of the capillary channels created in the folded device, where a noticeable alteration in the paper structure was observed because of the applied force.

**Fig. 1.**
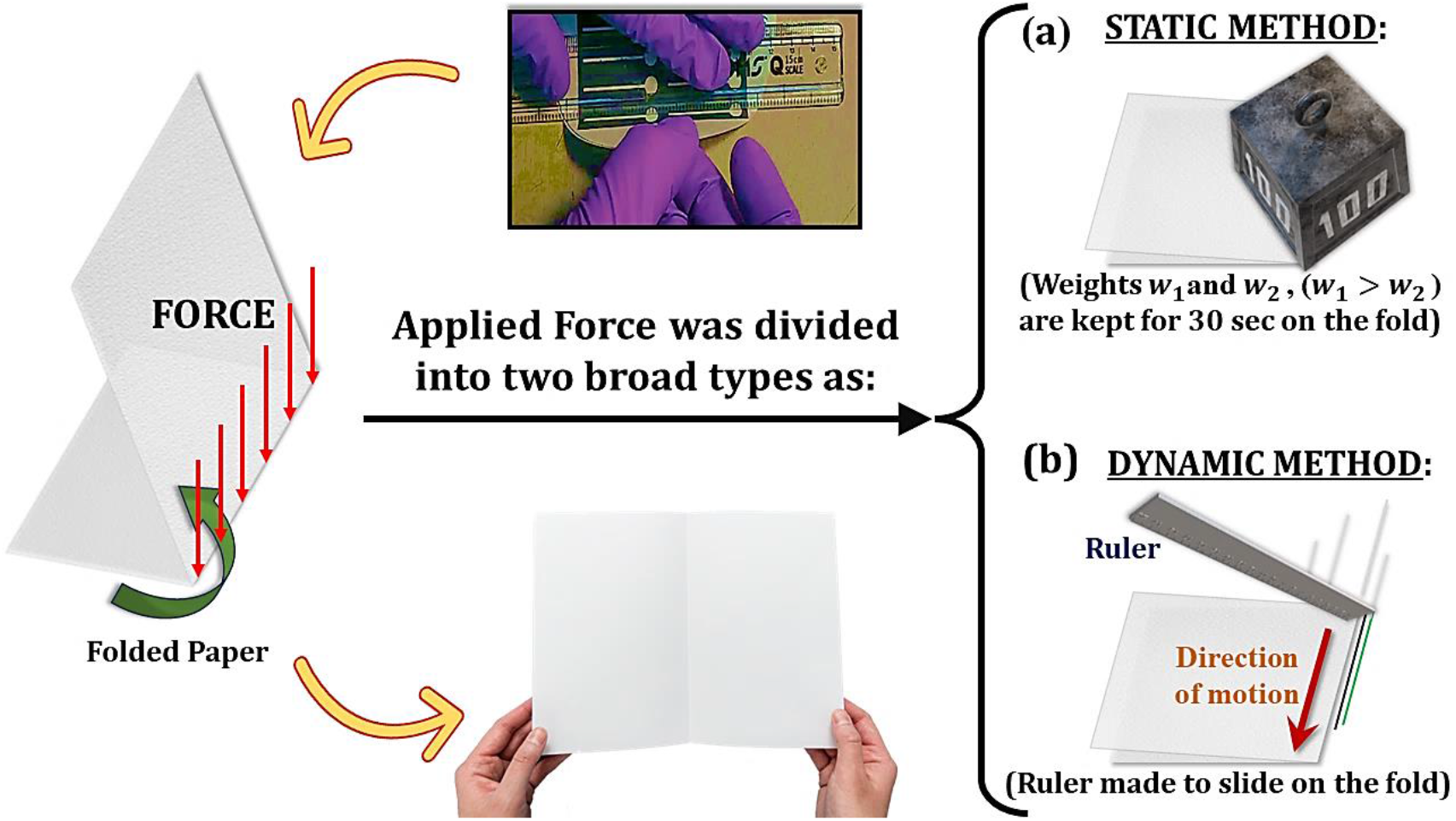
Schematic of the techniques employed to exert force to induce folding in paper channels. The force application can be categorised into **(a)** The Static Force Application Method, which involves the utilisation of two distinct fixed weights *w*_1_& *w*_2_, (*w*_1_ > *w*_2_), **(b)** The Dynamic Force Application Method, employing a ruler.

**Fig. 2.**
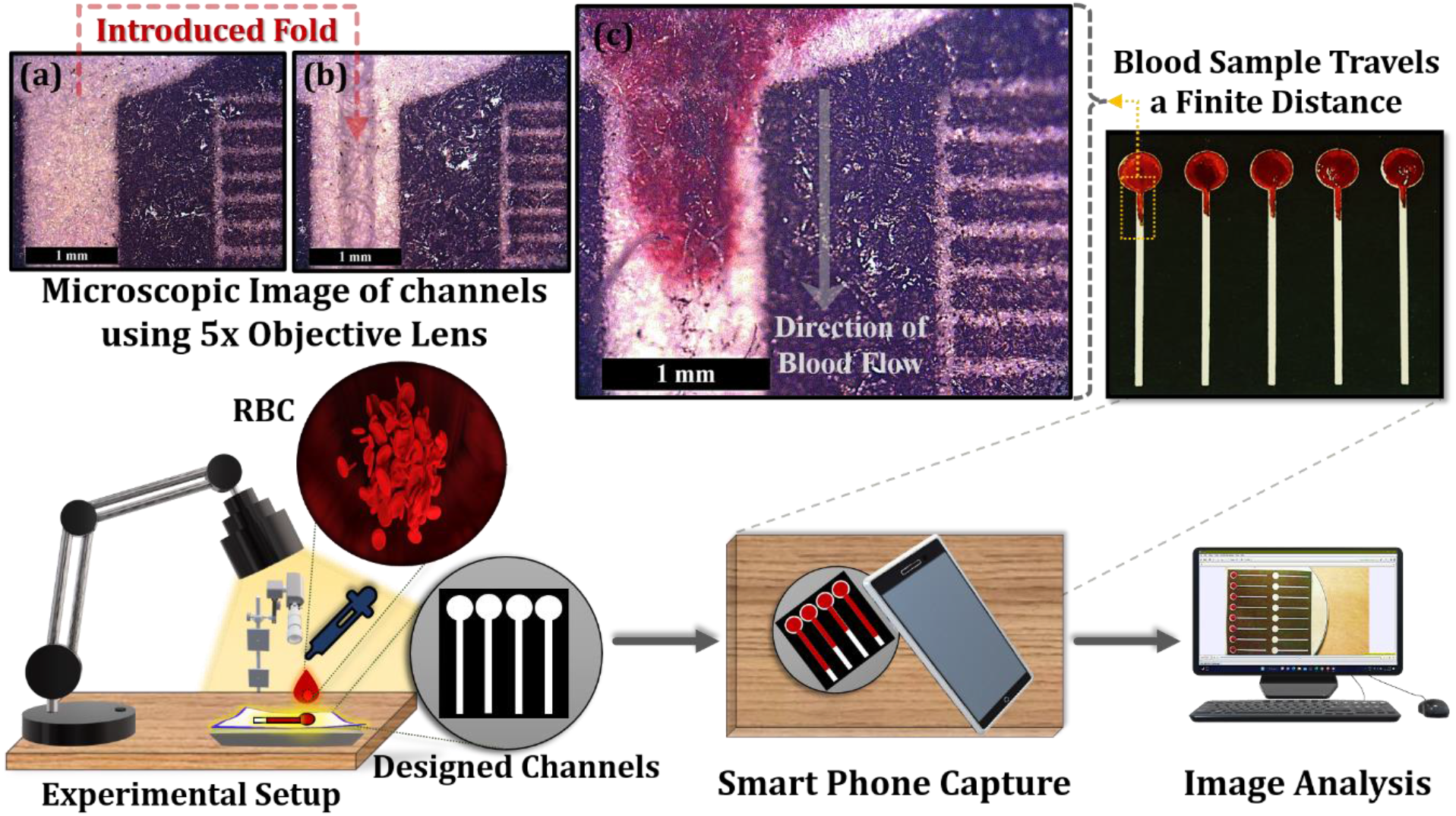
Experimental Setup employed for fluid front tracking using smartphone camera and subsequent analysis. The microscopic images of pattered channels on Grade-1 filter paper using a 5x Objective Lens **(a)** without any fold, **(b)** with introduced fold, and **(c)** flow trajectory of the applied sample in the fold-assisted paper channels.

### 2.2. Evaluation of Flow Dynamics

In order to maximize the utilisation of filter paper and reduce the cost per channel, each filter paper (dia. 110 mm and thickness 0.18 cm, measured using Vernier calliper) was patterned and sheared into sixteen channel strips for subsequent analysis. The wicking behaviour was investigated by dispensing 0.005g/ml solution of food dye (Orange Red Powder IH 7802) for enhanced flow visualization. The fluid front was recorded using a smartphone camera (Redmi Note 5 Pro) for a duration of 90 seconds. The experiments were conducted under controlled laboratory condition, with the temperature maintained at 27 ± 3 °C and the relative humidity ranging from 45%-55%. The data obtained from the experiments were evaluated using the open-source application Tracker (version 6.1.0).

### 2.3. Hematocrit Detection

#### 2.3.1. Baseline Calibration

For the second set of experiments, blood samples were collected from 7 healthy donors (refer **ESI. Table 1:** and **ESI. Table 2:** for the specifications) aged 20 to 60 years from B C Roy Technology Hospital, Indian Institute of Technology Kharagpur, India through venepuncture in EDTA tubes in accordance to standard laboratory protocol and after obtaining institutional ethical clearance (No. IIT/SRIC/DEAN/2022), ensuring participant’s confidentiality and informed consent. All the samples were used within 24 hrs of collection.

To establish a baseline calibration for distance travelled in paper for varied hematocrit contents, the supernatant plasma and buffy coat were eliminated from each sample. The obtained samples were centrifuged at 3000 rpm for 10 min using Tarson Spin win MC-01. The RBCs were then diluted using a phosphate buffer to achieve different cell concentrations ranging from 5% to 50%, with increments of 5%. The Phosphate buffer was utilised for dilution purposes to eliminate the potential influence of other CBC (Complete Blood Count) parameters, including proteins, ions, white blood cells (WBCs), and platelets, on flow dynamics. The samples (10 μL each) were spotted onto 10 simultaneous devices, and the mean and standard deviation of the final distance after 5 minutes were recorded (10 devices per conc. × 7 donors = 70 tests per conc.). The schematic of the setup and data acquisition procedure is shown in **Fig. 2**.

#### 2.3.2. Validation with Real Samples

The preliminary investigations for hematocrit measurement in the developed fast channels were conducted using 71 patients’ whole blood samples without any sample treatment. The final lengths traversed were recorded and analysed. The device was further validated by performing 54 blind tests of human patient samples based on the training data obtained from the previous two data sets. The blood samples were obtained from the Institute of Haematology and Transfusion Medicine, Calcutta Medical College, Kolkata using the previously discussed sample collection protocol from a pool of volunteers representing a diverse range of age and gender. For the above-mentioned tests, 10μL of blood samples were spotted onto simultaneous paper devices without any treatment or dilution and the distance travelled was recorded after 5 mins. The length travelled was measured from which the hematocrit range was determined. The obtained hematocrit range was then compared with the gold standard laboratory measurement data obtained from Automated Haematology Analyser (Sysmex KX-21). All the assays were performed without any controlled environmental condition to induce best reproducibility in POC assay. (The details of all blind test patient data have been added in the Supporting Information **ESI. 7**.)

## 3. Result and Discussion

### 3.1. Fold and Fast Flow Dynamics

The insights into the flow behaviour of the fabricated fast-channels can be obtained by analysing the temporal variation in the wicking length. The flow characteristics of the channels with folds are illustrated in **Fig.3(a)** and **Fig.3(b)**. The figures clearly outline a substantial increase in the wicking rate of channels with folds (≥ 20 cm in 90 seconds) as compared to the non-folded channels (≤ 10 cm in 90 seconds) that have identical patterned designs. The design dimensions were optimised for the inlet diameter (6 mm), channel width (1 mm) and sample volume (10 *μ*L). The wicking characteristics, upon varying these parameters, are obtained and the justification for selecting the optimised values is given in **ESI. Fig. 2**. and **ESI. Table 3**. The inlet section in this design is directly patterned onto the filter paper, in contrast to many previously published works utilizing a reservoir arrangement, for suitability in on field utilization.

The enhanced capillary wicking can be ascribed to the increased-average pore size of the folded crease formed under the action of the folding force, exerted along the designated axis. The observed fold in this context exhibits resemblance to bending followed by a buckle fold [22], wherein the paper fibres on the intrados will experience compression, while those on the extrados will undergo stretching, with maximum at the hinge points [23]. The non-stretch ability constraint of paper in the extrados will give rise to internal fibre tension, which gets conserved by tangential longitudinal strain along the machine direction (MD), coupled with the extrinsic curvature of the paper around the fold. The cross-sectional view of the fold and its SEM image, along with the top and bottom view of the stretched and compressed regions are shown in **Fig. 3**. The length of each channel is significantly greater as compared to the cross-section to induce plane strain condition throughout the folding process. The stress concentration may potentially induce a pore-size distribution along the thickness direction (ZD) of the paper, from minimum in the dense compressed region to maximum in the relaxed stretched region, see **ESI. Fig. 3**. As the fold develops, the neutral surface gradually approaches the inner arc boundary and the tensional features associated with the outer arc become increasingly prominent [24]. Consequently, the compressed region could get restricted to a narrow section, causing the larger pore regions to exert significant impact in determining the average pore radius along the flow direction. It is crucial to recognise that to create larger pores, it is required to apply an out-of-plane load instead of directly stretching the paper. This is because the anisotropic elastic constants of paper along the thickness direction (ZD) are around two orders of magnitude lower than the in-plane moduli [25], making it comparatively easier to deform the pores through folding.

**Fig. 3.**
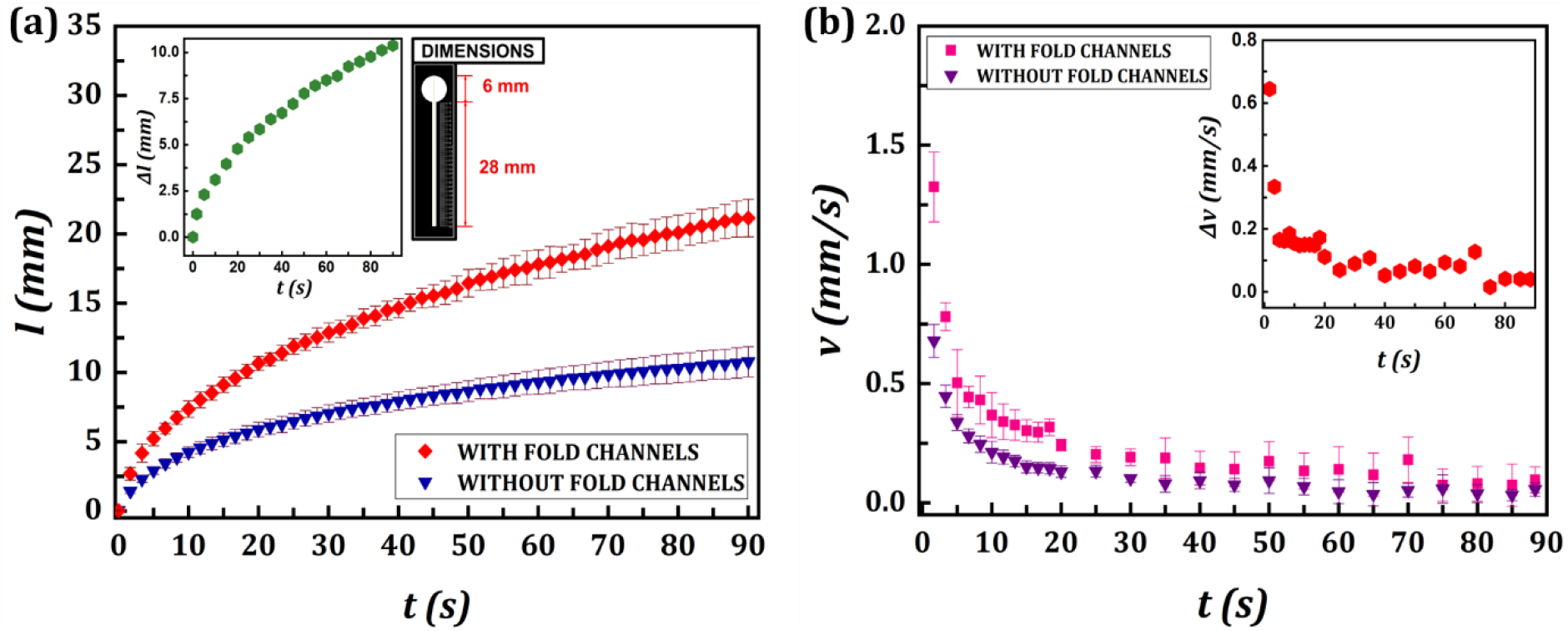
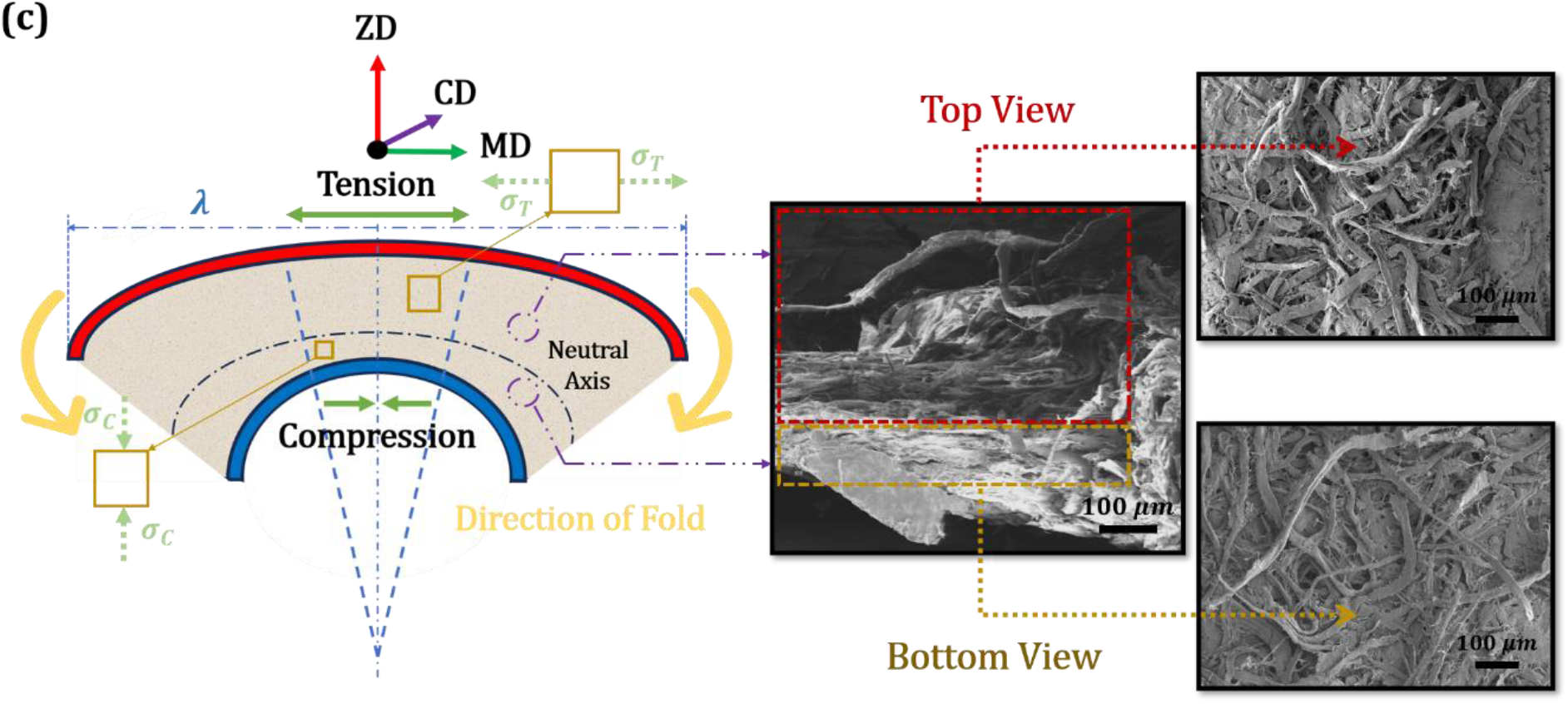
**(a)** Distance(mm) variation over time(s) for folded and non-folded channels with optimized parameters. The variation was repeated for 8 different channels and the average and standard deviations are plotted. **(b)** Velocity(mm/s) variation over time(s) is obtained from (a) using method of central difference. **(c)** Schematic and SEM Image of the fold cross-section, together with the top and the bottom views of the stretched (*σ*_*T*_) and compressed regions *(σ*_*C*_). (MD: Machine Direction, ZD: Thickness Direction & CD: Cross Direction)

Furthermore, considering the stochastic arrangement of the fibres in the paper matrix, the process of folding as a means of creating larger pores will effectively align them along the fold line, inducing a coordinated fibre arrangement. Hence, the exerted force fundamentally alters the inherent two-dimensional structure of the paper by generating fold creases that possess new geometrical configuration. This can be characterised by larger average pore size (***r***_***avg***_ ↑) and less tortuous (***τ*** ↓) porous network. These modifications facilitate improved capillary wicking within the folded channels. The observed difference in values of both the distance (Δ*l*(*t*)) and velocity (Δ*ν*(*t*)) measurements between folded and non-folded channels are also shown in **Fig.3(a)** and **Fig.3(b)** respectively. The trend clearly suggests that the rapid flow behaviour is consistently observed throughout the length and capillary wicking serves as the primary driving force for flow within the folded channels. As we can see, this technique can effectively accelerate even small liquid volumes (10 *μL*), establishing it as a potentially feasible platform for the investigation of biological fluids. Additionally, the presence of opposing body forces such as gravity do not alter the fast-flowing characteristics giving the flexibility to be utilized for assays in any orientation.

### 3.2. Methodology and Individuality Test

When the fracture energy (***G***_***F***_) of paper is considered from a macroscopic perspective, it is observed to be consistently proportional to the cross-sectional area **(*a***_***f***_) of broken inter-fibre bonds, **(*G***_***F***_ **∝ *a***_***f***_**) [**26]. This suggests that the breaking of these inter-fibre bonds will remain independent to any specific loading mode through which these bonds are broken. To substantiate this assertion and determine the most effective loading method for achieving consistent folding in our device, we utilised both Static and Dynamic force application strategies. The repeatability in the extent of fold formation through each technique was then examined by analysing their wicking characteristics. A significant fraction of bonding between paper fibres is mediated through inter-fibre contacts and the bond formation is governed by a complex interplay of several mechanisms. The intermolecular bonding interactions, including hydrogen bonding, electrostatic and van der Walls forces are only relevant for molecular contacts less than ∼300Å [27]. Similarly, the contribution from capillary bridging is constrained due to their confined occurrence to a rather small surface fraction without molecular contact. Therefore, the mechanical interlocking mechanism could dominate in establishing adhesion between the fibre surfaces at microscopic levels [28]. In conjunction to the rough surface topography, the inherent fibre diversity, morphological features such as pits and variations in residual strain from the pre-straining of top fibres could also contribute to the fibres being locked together mechanically. The results of our experiment, conducted using the static force method, showed that, by increasing the weight from *w*_2_to *w*_1_(*w*_1_ > *w*_2_), there is a notable improvement in the overall wicking rate, as depicted in **Fig. 4(a)**. This might be attributed to an increase in mechanical deformations inside the paper structure as the weight increases and thus inducing greater strains, see **ESI. Fig.4 (a) & (b)**. The mechanically entangled surfaces that interconnect the paper fibres could get more disrupted under a greater weight, causing the fibres to separate more and form bigger pores and contributing to an increased wicking rate. Furthermore, when subjected to the influence of a reduced load (*w*_2_), the wicking characteristics displays considerable variability in the wicking behaviour. This variation may potentially arise because of uneven dissipation of fracture energy when the fibres are subjected to deformation. This unregulated hysteresis observed in paper could be due to its highly anisotropic nature, preventing a sharp transition from elastic to elastic-plastic behaviour [26].

**Fig. 4.**
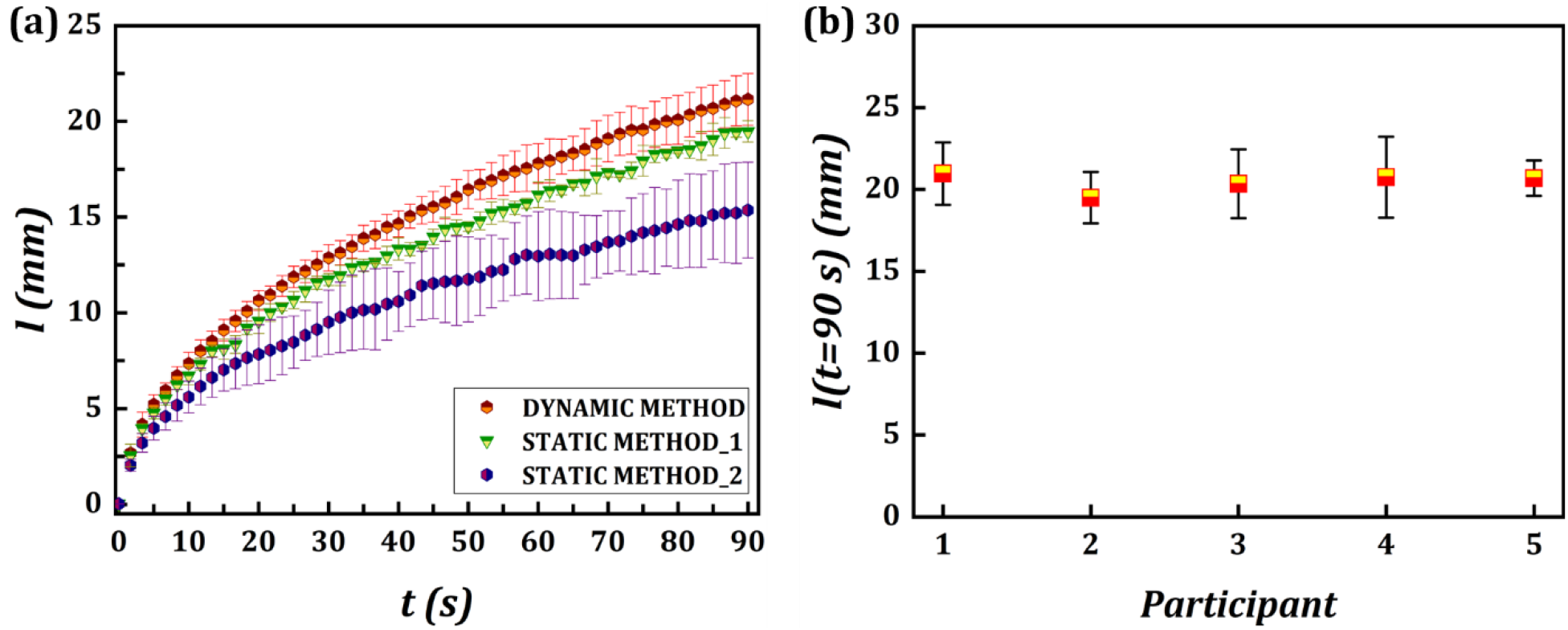
**(a)** Distance(mm) variation over time(s) for static and dynamic force application techniques. Static Method_1 (*w*_1_) and Static Method_2 (*w*_2_), **(b)** Individuality test results obtained from 5 different individuals. All the tests in this section are considered from making 8 folded channels using the dynamic method.

The Dynamic Method (inspired from origami) has been found to be the most favourable in terms of performance and production of uniform folds. Under the influence of an external force, the fibres initially undergo elastic deformation, which is governed by a power law model. As the applied force intensifies, the fibres undergo a critical transition to plastic deformation through the plastic hinge point, wherein the strain experienced by the fibres reaches maximum. Beyond this juncture, the paper fibres undergo negligible relaxation in response to additional forces [29]. This could be because the paper fibres would have reached the threshold to show substantial strain hardening in response to in-plane stresses along the MD [30]. The inability of paper fibres to revert their original shape after undergoing plastic deformation is coupled with their increased resistance to additional deformation owing to strain hardening. Therefore, the fibres in this maximum deformed condition will result in the formation of largest size pores. The utilisation of a ruler in this context facilitates the implementation of uniform and substantial force beyond this inflection point across the entire channel, thereby generating channels with maximum capillary rates. Although, same extent of force can also be applied using a static method, the choice of dynamic method is solely based on greater flexibility and lesser fabrication time (≤10 secs) while simultaneously ensuring precise control over the folding process. However, it is important to acknowledge that each approach possess its own advantages and trade-offs. While static approach offers better predictability, the dynamic approach presents superior adaptability and easy implementation in complex channel networks as in the present case.

To further establish the fabrication consistency and study the cross-individual variability of fold formation, the dynamic protocol was performed by five different participants on a set of eight allocated devices. The average distance covered by fluid during 90 secs, **(*l*(*t*** = **90 *s*)** across all users is ≈ 20.5 ± 1.8 mm, as depicted in **Fig. 4(b)**. This demonstrates that the use of ruler is an efficient and reliable technique for attaining precise and linear folds and the formation of fold can be consistent when created under identical conditions. In buckle fold formation, the ratio of fold wavelength to thickness, 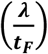 remain nearly constant, particularly in mechanically homogeneous materials and when formed under identical physical conditions [23]. Therefore, if both the requirements are met, the folds generated in paper will also be similar. Firstly, if consider the Weibull-modulus as a measure of uniformity of paper, typical values of which ranges from 10-15 [31]. This indicate that the paper can be regarded as a statistically uniform substrate. Furthermore, the tensile strength of paper, which significantly controls deformations in extrados areas, is mostly determined by the average macroscopic structure and the mean fibre properties [32]. Since it is not controlled by structural discontinuities and the structure of paper is statistically uniform, the tensile deformations in papers will also be relatively homogeneous. This implies that the initial criteria for achieving consistent folds is inherently satisfied. Secondly, beyond the plastic hinge point, as we discussed in the previous paragraph, the network mechanics plays a significant role in effectively transferring localised stress concentration to the adjacent fibres, making strain conditions in paper almost identical. So, if it is ensured that the critical force necessary to reach the plastic hinge point is applied by the dynamic method, the second criterion is also met. Therefore, any individual under normal loading will develop fold structures that are comparable (below initiating any fracture). The extent of fold formation, however, is heavily contingent on the paper type and its thickness, making it essential to consider those parameters a priori. If the fold-based devices were to be manufactured on a larger scale, the fabrication procedure can be improved using additional optimization and better equipment, such as Automated Buckle Folding machines, for more consistent fold and hence more consistent devices.

### 3.3. Model Development and Validation

In order to explain the imbibition characteristics through fold-introduced paper channels, a first-generation model has been developed elucidating the underlying physics of the flow. It is established in the study of buckle folds that there are negligible in-plane tensile strains beyond the creased zone, where these strains die-out rapidly [33]. Therefore, the flow domain can be divided into two distinct segments: the segment exhibiting a fold, characterised by a width of ***λ*** and the segment without fold, characterised by a width of **(*W*** − ***λ*)**, see **ESI. Fig. 5(a)**. Assuming the fluid properties (such as density, viscosity etc.) and prevailing flow conditions (laminar, fully developed 2-D flow) remain valid throughout the paper channel, the flow in each segment can be described by the simplified version of the Navier-Stokes equation. The velocity **(*ν*)** profile in the individual sections can thus be mathematically obtained using:

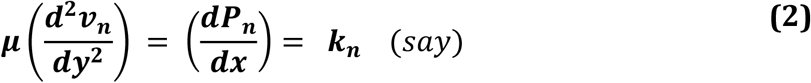

where *k*_*n*_ = *k*_*f*_ or *k*_*p*_, for flow within and outside the fold respectively, as the flow is driven by distinct pressure gradients 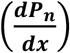, in the corresponding segments. Here, it is reasonable to assume a no-slip boundary condition for the fluid in contact with hydrophobic walls; and velocity and stress continuity conditions at the interface between fold and no-fold regions. Given the channel (width **(*W*)**/length **(*L*))** ∼ 0.03 − 0.1 to be small, the average pressure gradient (***k***_***avg***_) along the flow direction in terms of channel width **(*W*)** and fold wavelength **(*λ*)**, assuming an approximation of flow between parallel plates, can be derived using (**Eq. (2)**) as:

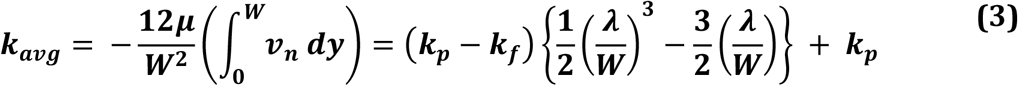

**Fig. 5.**
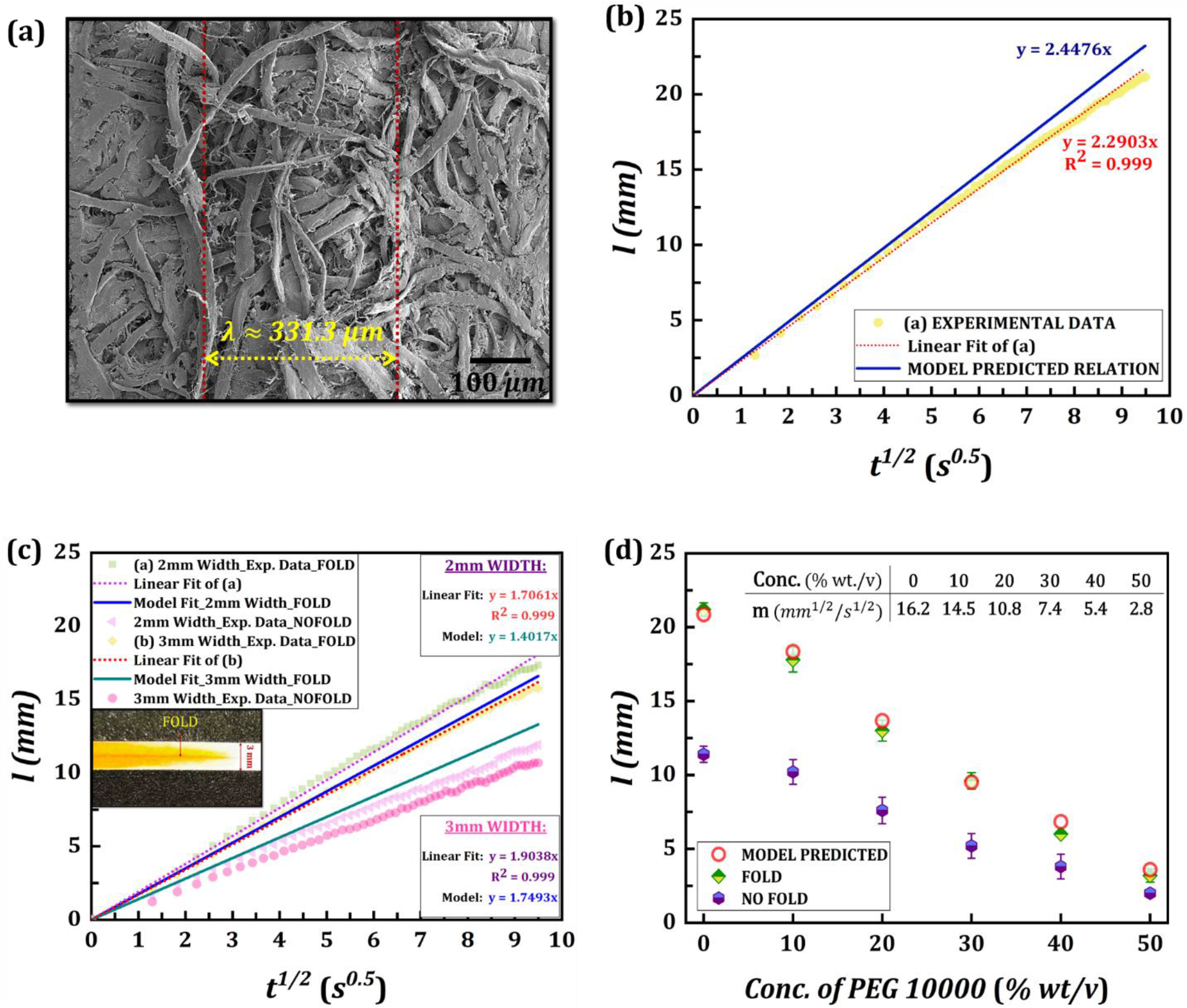
**(a)** SEM image of the channel cross-section where the fold is present. **(b)** Comparative representation of experimental and theoretically predicted wicking trend, for 1mm width channels **(c)** Deviation of the theoretical trend from the experimental data for 2 & 3mm width channels. The inset image represents the fluid trail for a 3mm width paper channel with a fold **(d)** Distance(mm) covered by PEG solution with different concentrations (% wt./v) in 90 secs (mean and SD of 5 replicates are plotted).

Furthermore, in accordance with the Hagen-Poiseuille equation, the fluid flow can be mathematically related to the characteristic dimension **(*l***_***c***_) of the channels, according to the relation: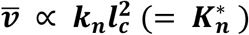. Thus, to approximate this flow behaviour in each segment of the folded channels, the pressure gradients **(*k***_***n***_) in (**Eq. (3)**) can be replaced with 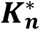. Considering that each characteristic section now consists of a network of cylindrical capillaries influenced by distinct driving forces, a mathematical relationship for the average pore radius (***r***_***avg***_) in terms of the established and measurable quantities can be devised by substituting *k*_*n*_ with the Laplace Pressure 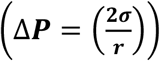. Hence, the expression for average pore radius through the channel can be expressed as:

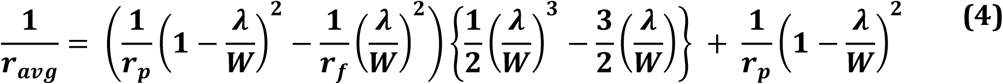

Here, ***r***_***f***_ and ***r***_***p***_ are the effective pore radius of the sections with fold and no-fold respectively. The estimated *λ* and *r*_*f*_ values for grade-1 filter paper (2*r*_*p*_ ≈ 11 *μm*) were obtained from analysing the SEM mages using ImageJ and were determined in the range of: *λ* ≈ 331 ± 20 *μm* and 2*r*_*f*_ ≈ 73 ± 11 *μm* at the hinge points, as shown in **Fig.5(a)**. The Lucas-Washburn equation for a liquid/vapor interface has undergone numerous tests and found to be generally valid throughout the range of *r* = 3 to 400 *μm* [34]. Thus, it can be extended to model the wicking behaviour through the creased paper channel by substituting (**Eq. (4)**) in (**Eq. (1)**) assuming homogeneous and isotropic pores. The trends shown in **ESI. Fig. 5(b)** illustrates the variation of the distance penetrated by fluid, (***l***(***t***)), with time (***t***) for both folded (*R*^2^ = 0.999) and unfolded (*R*^2^ = 0.998) paper channels according to (**Eq. (1)**). Since for both the channel types, the parameters influencing *m* ≈ 16.2 *mm*^1/2^/*s*^1/2^ (for food dye solution, see section 2.2) are presumed consistent, the difference in slope can be attributed to the difference in average pore radius 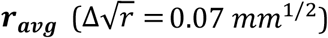 of the respective channels.

As shown in **Fig.5 (b)** for 1mm channel width, the theoretically predicted flow behaviour is in excellent agreement with the experimental results, with up to ∼ 93.1 % accuracy, demonstrating the viability of the proposed model. However, the model is observed to slightly overpredict the distance values associated with time, an outcome that is expected. This is because the governing equation is formulated based on certain set of assumptions, such as flow through cylindrical channels with rigid walls and no slip boundary conditions, which are not applicable for paper substrates. Additionally, parameters including porosity, tortuosity and effects like evaporation, non-specific absorption by paper fibres can also be included in the Lucas-Washburn equation, to improve the predictability of the model to estimate wicking in fold-based channels.

The flow rate is observed to decrease with increase in channel width **(*W*** ↑ ⇒ ***ν*** ↓) in case of both folded and unfolded channels, which align with the principles of conservation of mass. Additionally, as the width increases, the fraction of the region occupied by folds reduces 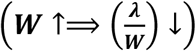, resulting in a flow rate approaching the flow rates observed in paper channels without folds. However, the proposed model equation tends to underpredict the distance variation with time as the channel width increases, the deviation becoming more prominent as the channel width is increased from 1mm (∼ 93.1 %) to 2mm (∼88.2 %) and subsequently to 3mm (∼66.8 %). This anomaly in fold introduced channels occur because the fluid flow concentrates in and within the proximity of the region subjected to folding, as highlighted in the image attached to **Fig. 5 (c)** for 3mm channel width. It is evident that while the fluid profile is parabolic in shape, the flow is predominantly towards the centre where the straight fold is present and does not cover the entire channel width. This occurs because the larger pore size within the folded section offers a less resistive pathway for rapid fluid flow and as a result instead of covering the entire channel width, the fluid selectively saturates the fold induced channel and its surroundings. At this point it diverges from the governing (**Eq. (3)**), which is formulated based on an average fluid velocity over the entire width of the flow channel. The fluid thus travels through a smaller effective area near the fold, rather than covering the entire channel width and is unimpeded by the hydrophobic walls in that region. Thus, the experimental wicking rate is observed to be higher (∼ 1.5 times for 3mm and ∼ 1.2 times for 2mm channel width) than that predicted from the model relation, which is also plotted in **Fig. 5 (c)**.

To further validate the model relation and verify the influence of bigger pores to induce rapid flow, the wicking experiments were conducted by varying different values of *m* = 16.2, 14.5, 10.8, 7.4, 5.4 and 2.8, where *m* is the fluid-specific component in the Lucas-Washburn equation that varies with contact angle, surface tension, and the fluid viscosity. Based on comparable blood rheology, PEG-10000 solutions with concentrations from 0-50% were considered to cover the viscosity range of healthy whole blood (∼3.5-5.5 cP [19]). Here, 0 % PEG solution correspond to DI water and for each PEG concentration, the tests were performed using 5 folded and non-folded channels. With the increase in PEG concentration, the value of *m* decreases due to increase in viscosity and surface tension, a characteristic established previously. The mean distance travelled by fluid in 90 secs and standard deviation, alongside values predicted from the model is plotted in **Fig. 5(d)**, clearly highlighting an average accuracy of ∼ 93.75 % of the proposed relation in predicting the wicking distance for different values of *m*.

### 3.4. Hematocrit Detection

The utilization of fold-induced paper channels offers advantages in terms of structural stability and the implementation of simplified printing techniques for device fabrication. This approach also simultaneously allows for the use of bigger pores to accelerate smaller liquid volumes, a critical aspect in point-of-care biofluid analysis. The efficacy of larger pores in facilitating rapid flow inside the developed paper channels is demonstrated through a length-based hematocrit sensor. The employment of distance-based approach in this context does not necessitate sophisticated instruments and on-site calibrators, rendering them easy to implement in field-settings. Furthermore, they demonstrate reduced susceptibility to human interference, enabling the provision of quantitative assessment with enhanced precision [35]. The use of fast-flowing paper channels for such distance-based measurement applications are advantageous, as ***l*(*t*)**_***fold***_ > ***l*(*t*)**_***unfold***_ ⇒ Δ***l*(*t*)**_***fold***_ > Δ***l*(*t*)**_***unfold***_ when Δ***t*** → **0**, indicating that these fast channels will result in greater change in length (Δ***l*(*t*)**_***fold***_) as compared to conventional paper channels (Δ***l*(*t*)**_***unfold***_) for specific variation in fluid properties **(*m*)**. This confers upon them a competitive edge to deliver superior performance at low-cost and operational time. The limit-of-detection (LOD) for these devices varies inversely with the difference in length **(*l*(*t*)**_***fold***_ − ***l*(*t*)**_***ufold***_) suggesting, the device will exhibit optimal accuracy when the fluid flow is completely halted **(*t*** → ∞) which is typically achieved in ∼5 min for blood samples.

With significant alteration in hematocrit levels, the blood flow through paper is affected, stemming from variation in blood viscosity and other pertinent hemorheological properties. As shown in **Fig. 6(a)**, the distance traversed by blood sample in the fabricated fast-channels patterned on grade-1 filter paper decreases with rise in hematocrit of blood **(% Hct** ↑ ⇒ ***l*** ↓), with marked difference in final length. This is anticipated because higher levels of hematocrit lead to an increase in blood viscosity, posing greater resistance to the flow of blood. Also, elevation in Hct levels influence its interaction with paper altering surface characteristics and capillary driving forces. In addition, a greater RBC fraction also contribute to inter-cellular interactions thereby leading to flow obstruction and reduced traverse length in the constricted capillaries of paper network. In contrast, flow through channels in grade-1 filter paper without induced fold showed minimal or insignificant blood flow. The flow is also observed to be negligibly small even when subjected to samples with hematocrit levels as low as 10 and 20%, **ESI. Fig. 6(a) & (b)**. The characteristic blood cell dimension typically ranges from ∼6−14*μm* and average pore sizes of grade-1 filter paper is ∼11 *μm*, thus the flow in no-fold paper channels is considerably impeded (Flow→ **0**) giving no distinction between various levels of hematocrit in blood. The use of grade-1 filter paper to perform this assay or in general to conduct any such similar assays is imperative due of its exceptional compatibility for using simplified fabrication techniques (such as laser printing) and hence aiding in the reduction of overall cost per device. Also, it exhibits a rather consistent distribution of pores and shows faster transfer rates that offer better analytical performance which can be utilized to overcome the limitations of µPADs [36].

**Fig. 6.**
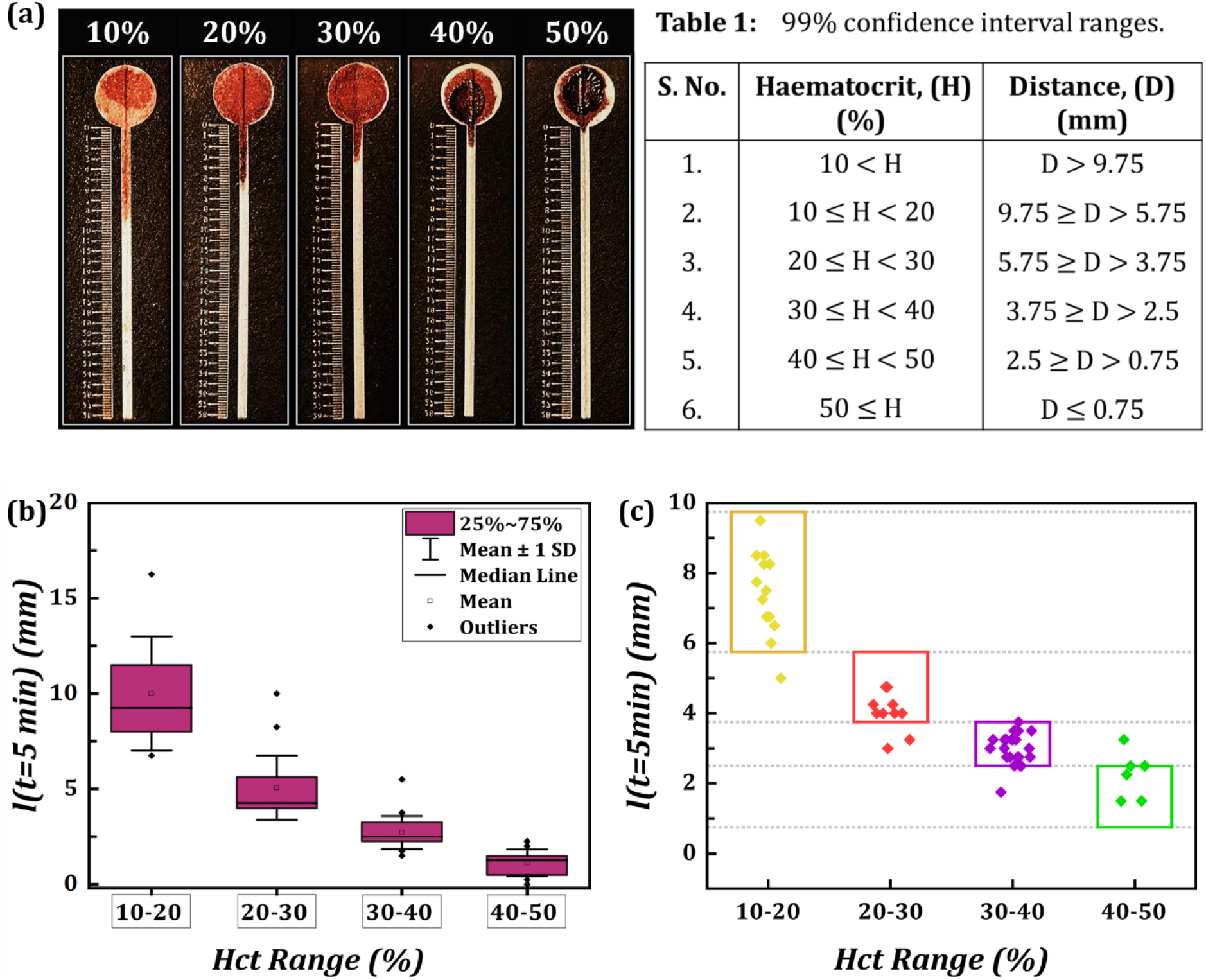
**(a)** The experimentally observed variations in the distances within the fold-induced paper channels for different % volume of RBCs (% Hct). **(b)** Results of comprehensive experimentation with the whole blood samples without any prior sample treatment or dilution. **(c)** The outcomes of blind test, with the coloured boxes indicating the length regions corresponding to the % Hct range and the horizonal lines represent the 99% confidence interval values from Table 1.

However, the tortuous network of paper and other CBC parameters could directly influence the blood flow through these folded channels, especially with whole blood without any sample preparation. This could pose a significant challenge in establishing a precise one-to-one correlation between hematocrit and length. Therefore, by combining length and Hct values in groups, a baseline calibration has been developed to determine length intervals within which a specific hematocrit level will vary. For a set of 7 healthy patient blood samples, a total of 7 × 10 × 10 = 700 tests for baseline calibration have been performed (see section 2.3.1). The experimental data obtained from the tests are given in **ESI. Table 4**. The calibrated length intervals were then determined using the 99% confidence interval equations **Eq. (5)** and **Eq. (6)**, given as:

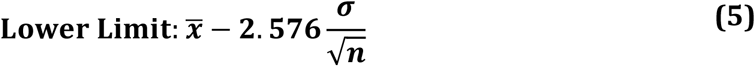

and

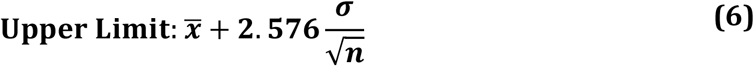

where 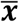 is the average distance travelled for known hematocrit %, ***σ*** is the standard deviation and ***n*** is the number of tests performed. The various parameters considered in the computation of 99% confidence interval is given in **ESI. Table 5**. and the obtained confidence interval of distance range in which the Hct levels will fall is tabulated in **Table: 1**. The interval data presented here clearly demonstrates notable differences in length for varied Hct levels. Thus, this characteristic can be effectively utilized as a differentiating parameter for determining the hematocrit range for an unknown blood sample.

In order to conduct a more comprehensive analysis of the detection capabilities, tests were performed to determine the hematocrit range of whole blood samples without any dilution or sample preparation. The tests repeated using 71 samples (data given in **ESI. Table 6**.) were able to generate reproducible ranges of length that align with the previously obtained calibration data, without any overlap, as shown in **Fig. 6(c)**. This indicates that significant change in hematocrit levels contribute substantially to its rheological behaviour to show observable length difference in fold-induced paper channels. There are other factors that may modulate blood rheology, such as plasma viscosity, shear rate, and blood cells characteristics, including deformability to show length difference in paper. But these factors could in general be significantly influential for individuals having certain medical conditions and disorders. However, such cases are not considered in the present analysis as per the individual’s medical report and to the best of our knowledge.

Additionally, a series of blind tests has also been performed from a group of both healthy and anaemic samples (selected based on their Hct levels) to rigorously evaluate the hematocrit detection capability of the fast-flowing sensor without any bias. The results of this extensive blind test, as shown in **Fig. 6(d)** demonstrate that the measured ranges of hematocrit using our paper devices are in good consistency with the reference values obtained from the commercial haematocytometer. The data is provided in **ESI. Table 7**. From a total of 54 samples chosen at random and subjected to analysis, the device showed a sensitivity of 90.74% underscoring the precision and versatility in accurately measuring hematocrit levels across a broad spectrum of blood samples (the images of some of the blind tests is given in **ESI. Table 8**). Sensitivity in this context refers to the ability of the test to correctly identify true positives and the outcome indicates a strong correlation between the test and the target condition, thereby affirming the utility and accuracy of our testing approach. Therefore, the device has potential as a viable platform for preliminary screening of anaemic patients in resource limited settings. The blood sample test findings presented in this section further contribute to our understanding of the process of larger pore formation in folded paper channels. It opens the possibility for including other micrometre-sized entities, such as cells, microbeads, and bacteria, as test components. This underscores the clinical relevance and reliability of these types of measurements for biomedical applications in point-of-care diagnostics.

## 4. Conclusion

In this work, we have proposed a novel and efficient paper-microfluidic platform using grade-1 filter-paper that enables rapid fluid-flow (∼2X) by employing a single straight fold. The folds were implemented via a ruler in a significantly reduced timeframe (≤ 10 s), while the devices were fabricated using a table-top laser printer at a very low expense without any complexity. The developed fast-channels employed to determine the blood Hct levels could precisely differentiate the ranges with 90.74% sensitivity when validated with a commercial haematocytometer. Prior to sensing, the design was optimized and the channels could effectively accelerate even smaller liquid volumes (10*μ*L) providing a distinct advantage for manipulating biofluids. A first-generation Washburn form model with an adjusted estimation of *r*_***avg***_ is proposed, accurately predicting the fast-flow characteristics (∼93.1%) and thereby explaining the underlying physics. But there is a limit to how much folding can increase the pore size and augment the flow rates. Thus, multiple folds and different filter paper grades could be explored in conjunction with other factors for achieving desired flow without compromising the structural stability of µPADs. The fast paper device exhibits promising capabilities to develop medical devices with complex networks and faster response, improving healthcare access and rapid detection in the near future.

## CRediT authorship contribution statement

**Amaan Dash**: Conceptualization, Data curation, Formal analysis, Investigation, Visualization, Methodology, Writing – Original draft; **Manikuntala Mukhopadhyay**: Conceptualization, Formal analysis, Investigation, Writing – review & editing; **Jyoti Shaw**: Data curation, Investigation, Validation; **Maitreyee Bhattacharya**: Supervision, Investigation, Validation; **Sunando DasGupta**: Conceptualization, Investigation, Project administration, Funding acquisition, Supervision, Writing – review & editing

## Acknowledgements

The authors gratefully acknowledge the financial support provided by the Indian Institute of Technology Kharagpur, India [Sanction Letter no.: IIT/SRIC/ATDC/CEM/2013-14/118, dated 19.12.2013]. AD is thankful to the Ministry of Education, Government of India and IIT Kharagpur for the Prime Minister’s Research Fellowship (PMRF). We acknowledge the co-operation of Pathology division of the B. C. Roy Technology Hospital, Indian Institute of Technology Kharagpur, India, and Institute of Haematology & Transfusion Medicine, Medical College, Kolkata, India for providing blood samples in accordance with the Institute’s Ethical Guidelines as well as for providing gold standard hematology analysis data.

