## Supplemental File for "RAPID HEMATOCRIT ESTIMATION USING A FOLD-CREASE INDUCED FAST FLOWING PAPER SENSOR"

### ESI. 1. Device Fabrication:

The channels were designed using open-source application Inkscape (1.1.2). The printed devices were then placed on a hot plate for 30 min. A circular steel block of diameter 110 mm and thickness 27mm, preheated to 180°C was placed on top of it to ensure uniform ink penetration from both sides and form leakage free hydrophobic barriers.[1]

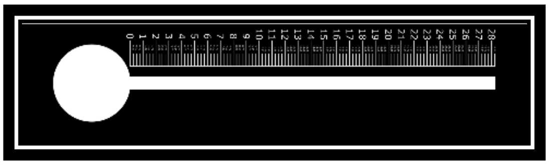

### ESI. 2. Healthy Blood Specifications:

Individuals, with $50 \pm 20$ kg of body-weight, $73 \pm 4$ beats/min of resting heart rate, $120.5 \pm 7$ mmHg of systolic and $84 \pm 3$ mm Hg of diastolic blood pressure including non-smokers and devoid of any disease and disorder are included in the analysis.

**ESI. Table 1:** The specific blood parameters and their ranges obtained from Automatic Haematology Analyser (Sysmex, KX-21), for the healthy individuals, are listed below.

| SR. NO. | Parameter | Range | Unit |
|---|---|---|---|
| 1 | WBC | 3.5-9.5 | $10^3$/μL |
| 2 | Neu% | 40-75 | % |
| 3 | Lym% | 20-50 | % |
| 4 | Mon% | 3-10 | % |
| 5 | Eos% | 0.4-8 | % |
| 6 | Bas% | 0-1 | % |
| 7 | Neu# | 1.8-6.3 | $10^3$/μL |
| 8 | Lym# | 1.1-3.2 | $10^3$/μL |
| 9 | Mon# | 0.1-0.6 | $10^3$/μL |
| 10 | Eos# | 0.02-0.52 | $10^3$/μL |
| 11 | Bas# | 0-0.06 | $10^3$/μL |
| 12 | ALY# | 0-0.2 | $10^3$/μL |
| 13 | ALY% | 0-2 | % |
| 14 | LIC# | 0-0.02 | $10^3$/μL |
| 15 | LIC% | 0-0.1 | % |
| 16 | NRBC | 0.00 | $10^3$/μL |
| 17 | NRBC% | 0.00 | % |
| 18 | RBC | 3.8-5.8 | $10^6$/μL |
| 19 | HGB | 11-17 | g/dL |
| 20 | HCT | 35-50 | % |
| 21 | MCV | 82-100 | fL |
| 22 | MCH | 27-34 | Pg |
| 23 | MCHC | 31-35 | g/dL |
| 24 | RDW-CV | 11-16 | % |
| 25 | RDW-SD | 35-56 | fL |
| 26 | PLT | 125-350 | $10^3$/μL |
| 27 | MPV | 6.5-12 | fL |
| 28 | PDW-SD | 9-17 | fL |
| 29 | PDW-CV | 10-17.9 | % |
| 30 | PCT | 0.1-0.2 | % |
| 31 | P-LCR | 11-45 | % |
| 32 | P-LCC | 30-90 | $10^3$/μL |

**ESI. Table 2:** Hematocrit data of 7 selected heathy samples for baseline calibration.

| **SR.NO.** | 1 | 2 | 3 | 4 | 5 | 6 | 7 |
|---|---|---|---|---|---|---|---|
| **HCT (%)** | 38.1 | 38.4 | 43.0 | 40.5 | 40.8 | 48.4 | 43.4 |

### ESI. 3. Device Optimization:

**ESI. Table 3:** Optimised parameters for fast flowing folded channels.

| SR. No. | Parameters | Selected | Justification |
|---|---|---|---|
| 1. | Inlet Diameter **(ESI. Fig. 1(a))** | 6 mm | No significant back flow (unlike 7mm), Less toner cost as compared to 5mm. (**ESI. Fig. 2(a)**) |
| 2. | Channel Width **(ESI. Fig. 1(b))** | 1mm | As $W\uparrow \Longrightarrow$ Flow $\downarrow$, justified by continuity eq. and $(\lambda/W) \rightarrow 0$ with $\uparrow$ in $W$. (**ESI. Fig. 2(b)**) |
| 3. | Sample Volume **(ESI. Fig. 1(c))** | 10 µL | No wicking improvement above 10µL due to greater intermolecular attraction (Cohesive Forces > Adhesive Forces). (**ESI. Fig. 2(c)**) [2] |
| 4. | Gravity Effect | NO* | Small sample volume. **(ESI. Fig. 2(d))** |
| 5. | Evaporation Effect | NO* | No observable difference (Scotch Magic Tape, 3M India Ltd.), Microcapillaries formed by uneven tape-paper contact increasing wicking variation. **(ESI. Fig. 2(e) & 2(f))** |

*NO: Not Observed (no effect of this parameter is recorded)

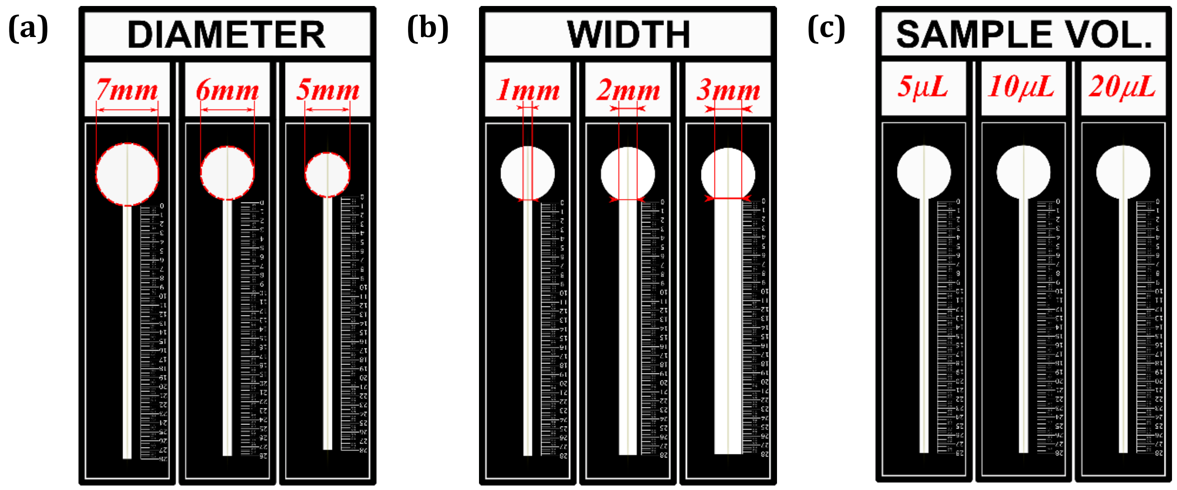


**ESI. Fig. 1** All the design parameters of the fast-flowing paper channel that are optimised. **(a)** The inlet diameter section of the channel was varied between 5, 6 and 7mm. **(b)** The channel width was varied between 1, 2 and 3mm., **(c)** The sample volume was varied from 5, 10 and 20 $\mu L$. Length of each channel was kept constant at 28 mm.

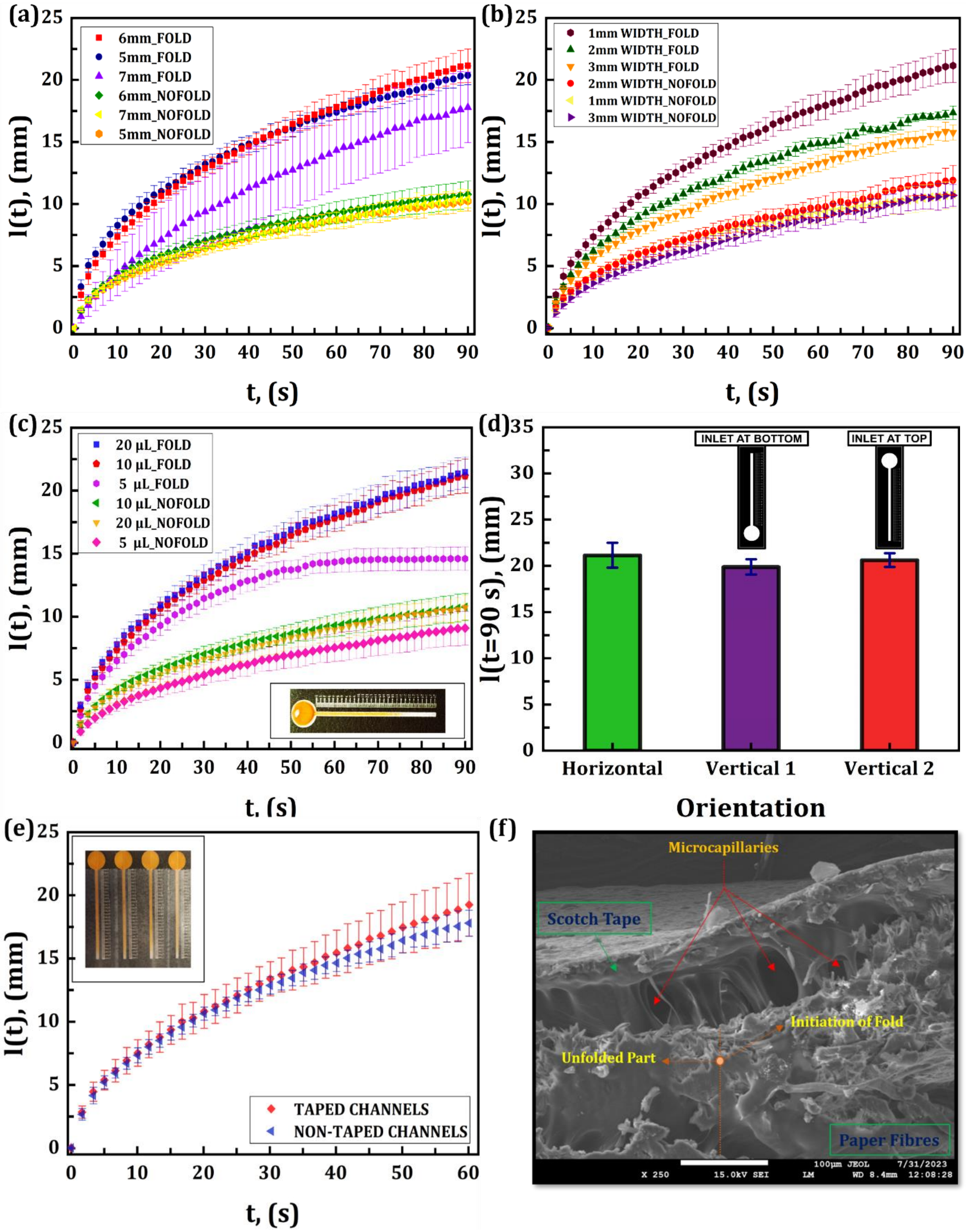


**ESI. Fig. 2** **(a)**, **(b)** & **(c)** Distance(mm) variation over time(s) for folded and non-folded channels for inlet section diameter, channel width and sample volume optimization respectively. **(d)** Effect of gravity in different orientations. **(e)** Distance(mm) variation over time(s) for taped and non-taped channels **(f)** Cross-Sectional view of the contact between Scotch Tape and paper fibres, along with formation of microcapillaries. All the graphs were obtained by repeating the characteristics for 8 different channels and the average and standard deviations are plotted.

**ESI. 4. Fast Flow Dynamics:**

SEM images are used to illustrate the contrasting pore-size difference between the folded-creased regions, at the fold hinge points and the non-folded paper network. The images demonstrate that folding causes the interconnecting fibres to relax, resulting in the existing pores to become larger.

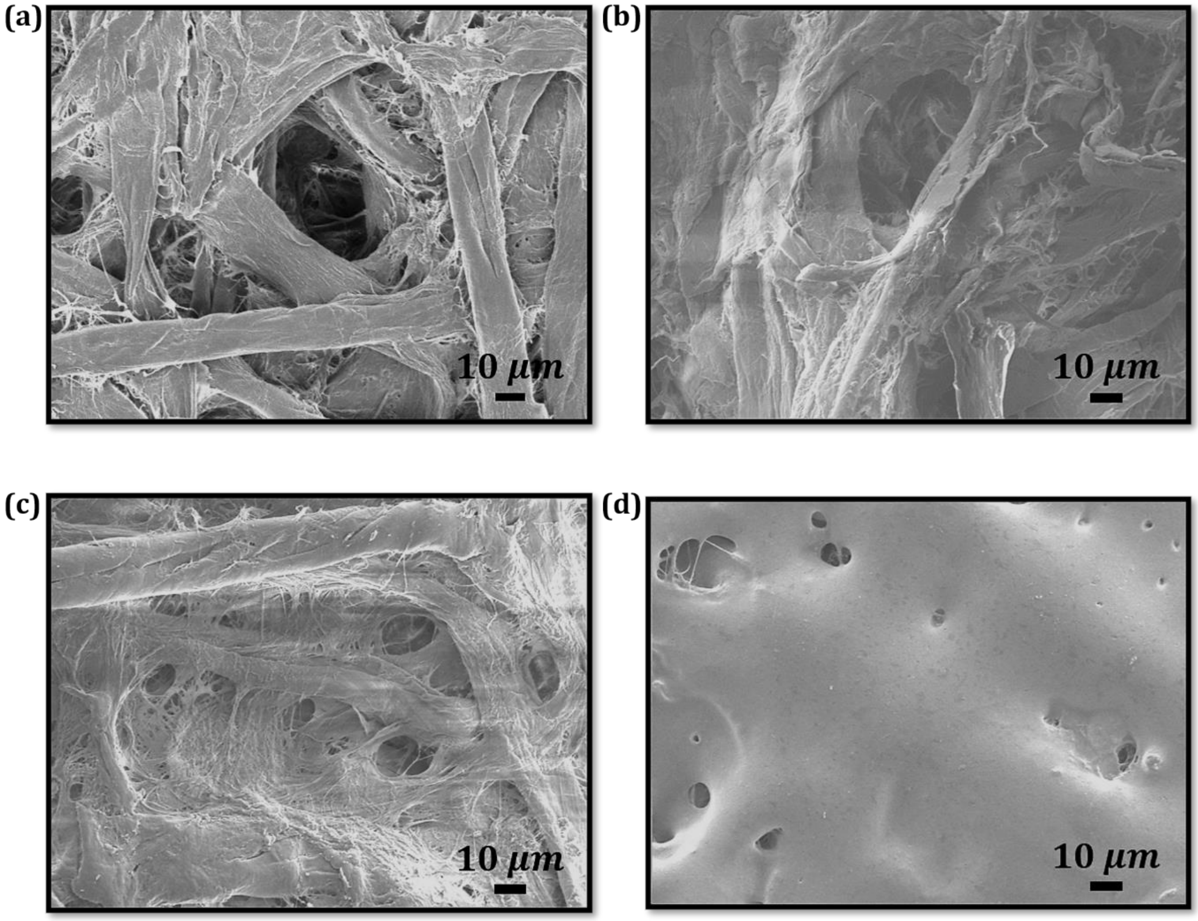


**ESI. Fig. 3** **(a)** & **(b)** SEM Images of hinge points of fold-introduced paper channels for two different devices on a grade-1 filter paper. **(c)** SEM image of grade-1 filter paper **(d)** SEM Image of laser printed hydrophobic regions.

**ESI. 5. Methodology and Individuality Test:**

SEM Images of the folded-creased region under two distinct loads, applied using the static force application strategy. It is evident from the images that as the weight increases, the fibres exhibit a greater tendency to distort. A further observation of interest is the consistent similarity in the dimension of the fold wavelengths across all the various techniques used, suggesting a high degree of consistency in the production of folds.

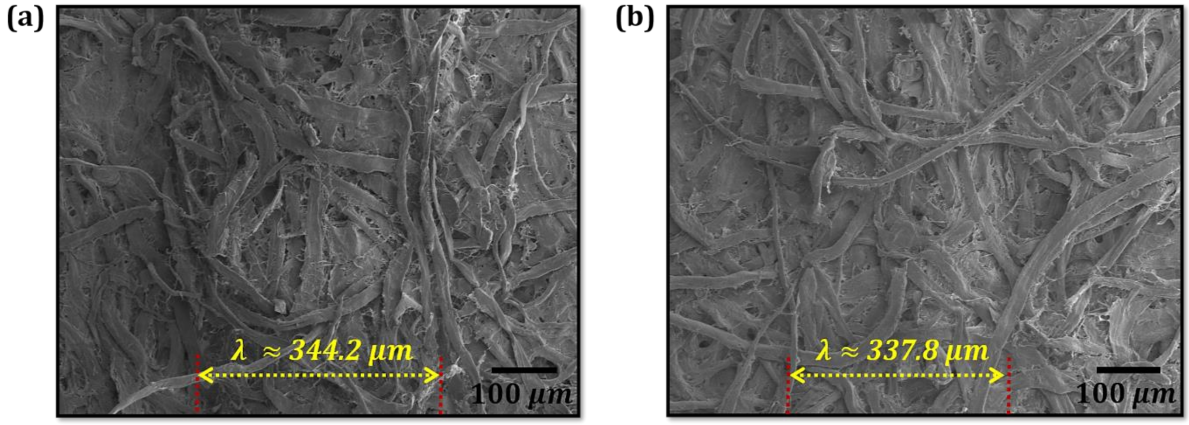


**ESI. Fig. 4** **(a)** & **(b)** SEM Images of the extent of deformation using weight $w_2$ and $w_1$, where $(w_1 > w_2)$ respectively. The simultaneous fold wavelengths are also highlighted in each image.

### ESI. 6. Model Development and Validation:

The experimentally obtained wicking characteristics observed in both the channels with folds and the channels without folds align well with the Lucas-Washburn equation. This reaffirms our previously established fact that the driving force in the folded channel is identical to that of the standard paper channels, which is the capillary driving force.

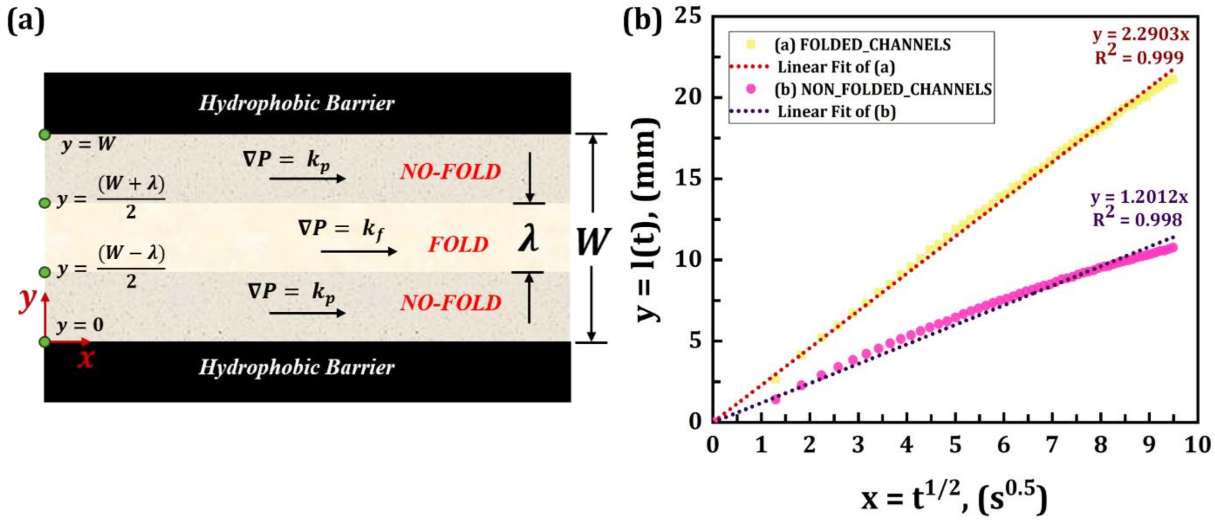


**ESI. Fig. 5** **(a)** Schematic of the fold-introduced paper channel used for model development. **(b)** Distance variation with square root of time ($t^{0.5}$), obtained from the experimental Distance (mm) Vs Time ($s^{0.5}$) data, for 1mm Channel Width.

### ESI. 7. Application:

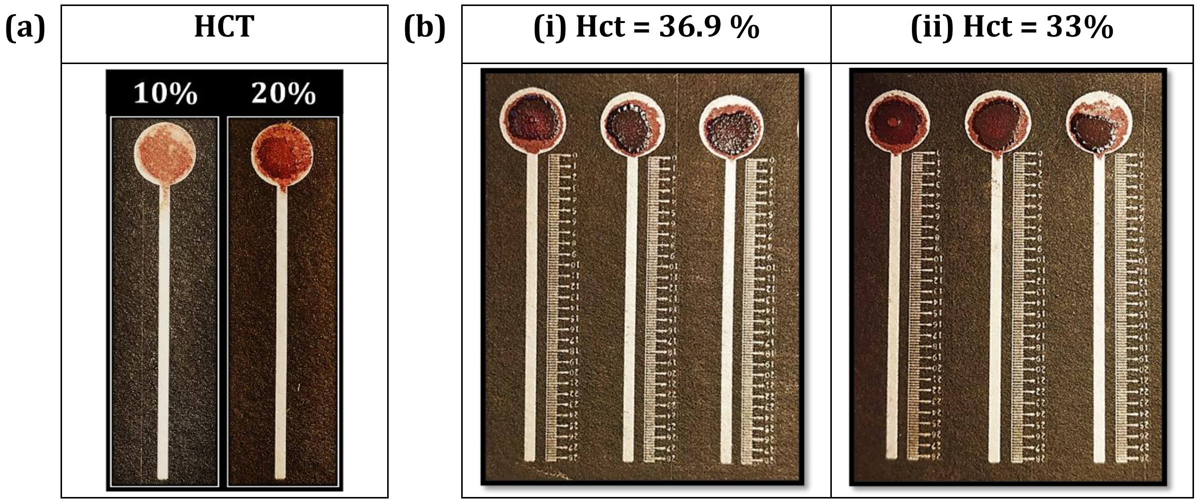


**ESI. Fig. 6** **(a)** Flow → 0 for 10% and 20% diluted Hct blood samples in non-folded Grade-1 filter paper. **(b)** Flow → 0 for (i) 36.9% & (ii) 33% blood Hct, without any dilution or sample treatment, on three simultaneous devices patterned on Grade-1 filter paper without any fold introduced.

**ESI. Table 4:** Data set obtained for 7 healthy blood samples. Each sample was diluted to 10 concentrations, from 5, 10…. upto 50 % Hct and then each concentration from every sample was dispensed onto 10 simultaneous paper devices, (7× 10=70 raw test results for each concentration). A total of 70 × 10 = 700 tests for baseline calibration. (The samples are acquired from B.C Roy Technology Hospital, Indian Institute of Technology, Kharagpur)

| SR. No. | % HEMATOCRIT | | | | | | | | | |
|---|---|---|---|---|---|---|---|---|---|---|
| | 5 | 10 | 15 | 20 | 25 | 30 | 35 | 40 | 45 | 50 |
| **1** | 19.5 | 12.75 | 15 | 9 | 13 | 6 | 4.5 | 7 | 2.75 | 2.75 |
| **2** | 16.75 | 12.5 | 13 | 8.25 | 11.75 | 5.25 | 4.5 | 3.5 | 2.75 | 2.5 |
| **3** | 16.25 | 12.5 | 13 | 8 | 11.75 | 4.75 | 4.25 | 3.25 | 2.75 | 1.75 |
| **4** | 15.5 | 11.75 | 12.25 | 7.75 | 11 | 4.5 | 4 | 3.25 | 2.5 | 1.5 |
| **5** | 15.25 | 11.75 | 11.5 | 7.75 | 9.25 | 4.5 | 4 | 3 | 2.5 | 1.5 |
| **6** | 15 | 11.5 | 10.75 | 7.75 | 9 | 4 | 3.75 | 2.75 | 2.25 | 1.5 |
| **7** | 14.75 | 11.5 | 10.25 | 7.5 | 9 | 4 | 3.75 | 2.75 | 2.25 | 1.5 |
| **8** | 14.25 | 11.5 | 9.75 | 7.5 | 8.75 | 4 | 3.75 | 2.75 | 2 | 1.5 |
| **9** | 13.25 | 11.5 | 9.75 | 7.25 | 8.25 | 4 | 3.75 | 2.75 | 2 | 1.25 |
| **10** | 13.25 | 11 | 8.75 | 7.25 | 7.75 | 4 | 3.5 | 2.75 | 2 | 1.25 |
| **11** | 13 | 11 | 8.75 | 6.75 | 7.25 | 4 | 3.5 | 2.75 | 2 | 1.25 |
| **12** | 12.75 | 11 | 8.75 | 6.75 | 6.75 | 3.75 | 3.25 | 2.75 | 2 | 1.25 |
| **13** | 12.5 | 10.75 | 8.75 | 6.75 | 6.5 | 3.75 | 3.25 | 2.75 | 2 | 1.25 |
| **14** | 12.5 | 10.5 | 8.75 | 6.5 | 6.25 | 3.75 | 3 | 2.75 | 2 | 1.25 |
| **15** | 12 | 10.25 | 8.25 | 6.5 | 6.25 | 3.75 | 3 | 2.75 | 2 | 1.25 |
| **16** | 12 | 10.25 | 8.25 | 6.5 | 6 | 3.75 | 3 | 2.5 | 1.75 | 1.25 |

| | | | | | | | | | | |
|---|---|---|---|---|---|---|---|---|---|---|
| **17** | 12 | 10 | 8.25 | 6 | 6 | 3.75 | 3 | 2.5 | 1.75 | 1.25 |
| **18** | 12 | 10 | 8.25 | 5.75 | 5.75 | 3.75 | 3 | 2.5 | 1.75 | 1 |
| **19** | 11.75 | 10 | 8 | 5.75 | 5.5 | 3.75 | 3 | 2.25 | 1.75 | 1 |
| **20** | 11.75 | 10 | 8 | 5.75 | 5.25 | 3.75 | 2.75 | 2.25 | 1.75 | 1 |
| **21** | 11.75 | 9.75 | 7.75 | 5.75 | 5.25 | 3.75 | 2.75 | 2.25 | 1.75 | 1 |
| **22** | 11.5 | 9.75 | 7.75 | 5.5 | 5.25 | 3.75 | 2.75 | 2.25 | 1.75 | 1 |
| **23** | 11.5 | 9.75 | 7.5 | 5.5 | 5.25 | 3.75 | 2.75 | 2.25 | 1.75 | 1 |
| **24** | 11.25 | 9.75 | 7.5 | 5.5 | 5 | 3.75 | 2.75 | 2.25 | 1.75 | 1 |
| **25** | 11.25 | 9.75 | 7.25 | 5.5 | 5 | 3.75 | 2.75 | 2.25 | 1.5 | 1 |
| **26** | 11.25 | 9.75 | 7.25 | 5.5 | 5 | 3.75 | 2.75 | 2.25 | 1.5 | 1 |
| **27** | 11.25 | 9.75 | 7 | 5.5 | 5 | 3.75 | 2.75 | 2.25 | 1.5 | 1 |
| **28** | 11.25 | 9.5 | 7 | 5.25 | 4.75 | 3.75 | 2.75 | 2.25 | 1.5 | 1 |
| **29** | 11 | 9.5 | 6.75 | 5.25 | 4.75 | 3.5 | 2.75 | 2.25 | 1.5 | 1 |
| **30** | 11 | 9.25 | 6.75 | 5.25 | 4.75 | 3.5 | 2.75 | 2.25 | 1.5 | 1 |
| **31** | 11 | 9.25 | 6.75 | 5.25 | 4.75 | 3.5 | 2.75 | 2.25 | 1.5 | 0.75 |
| **32** | 11 | 9.25 | 6.75 | 5 | 4.5 | 3.5 | 2.5 | 2.25 | 1.5 | 0.75 |
| **33** | 11 | 9 | 6.5 | 5 | 4.25 | 3.25 | 2.5 | 2.25 | 1.5 | 0.75 |
| **34** | 10.75 | 9 | 6.25 | 5 | 4.25 | 3.25 | 2.5 | 2 | 1.5 | 0.75 |
| **35** | 10.75 | 9 | 6.25 | 4.75 | 4.25 | 3.25 | 2.5 | 2 | 1.5 | 0.75 |
| **36** | 10.75 | 8.75 | 6.25 | 4.75 | 4.25 | 3.25 | 2.5 | 2 | 1.5 | 0.75 |
| **37** | 10.5 | 8.75 | 6.25 | 4.75 | 4.25 | 3.25 | 2.5 | 2 | 1.5 | 0.75 |
| **38** | 10.5 | 8.75 | 6 | 4.75 | 4.25 | 3.25 | 2.5 | 2 | 1.5 | 0.75 |
| **39** | 10.5 | 8.75 | 6 | 4.5 | 4.25 | 3.25 | 2.5 | 2 | 1.5 | 0.75 |
| **40** | 10.5 | 8.75 | 6 | 4.5 | 4.25 | 3.25 | 2.5 | 2 | 1.5 | 0.75 |
| **41** | 10.5 | 8.75 | 6 | 4.5 | 4.25 | 3.25 | 2.5 | 2 | 1.5 | 0.75 |
| **42** | 10.5 | 8.75 | 5.75 | 4.5 | 4 | 3.25 | 2.25 | 1.75 | 1.25 | 0.75 |
| **43** | 10.5 | 8.5 | 5.75 | 4.5 | 4 | 3.25 | 2.25 | 1.75 | 1.25 | 0.5 |
| **44** | 10.25 | 8.5 | 5.75 | 4.25 | 4 | 3.25 | 2.25 | 1.75 | 1.25 | 0.5 |
| **45** | 10.25 | 8.5 | 5.75 | 4.25 | 4 | 3 | 2.25 | 1.75 | 1.25 | 0.5 |
| **46** | 10 | 8.5 | 5.75 | 4.25 | 4 | 3 | 2.25 | 1.75 | 1.25 | 0.5 |
| **47** | 10 | 8.25 | 5.75 | 4.25 | 4 | 3 | 2.25 | 1.75 | 1.25 | 0.5 |
| **48** | 10 | 8.25 | 5.75 | 4.25 | 4 | 3 | 2.25 | 1.75 | 1.25 | 0.5 |
| **49** | 10 | 8.25 | 5.5 | 4.25 | 4 | 3 | 2.25 | 1.75 | 1.25 | 0.5 |
| **50** | 10 | 8.25 | 5.5 | 4.25 | 4 | 3 | 2.25 | 1.75 | 1.25 | 0.5 |
| **51** | 10 | 8 | 5.5 | 4.25 | 3.75 | 3 | 2 | 1.75 | 1.25 | 0.5 |
| **52** | 10 | 7.75 | 5.5 | 4.25 | 3.75 | 3 | 2 | 1.5 | 1 | 0.5 |
| **53** | 10 | 7.75 | 5.5 | 4 | 3.75 | 3 | 2 | 1.5 | 1 | 0.5 |
| **54** | 10 | 7.75 | 5.5 | 4 | 3.75 | 3 | 2 | 1.5 | 1 | 0.5 |
| **55** | 10 | 7.75 | 5.5 | 4 | 3.75 | 3 | 2 | 1.5 | 1 | 0.5 |
| **56** | 9.75 | 7.75 | 5.25 | 4 | 3.75 | 3 | 2 | 1.5 | 1 | 0.5 |
| **57** | 9.75 | 7.5 | 5.25 | 4 | 3.5 | 3 | 2 | 1.5 | 1 | 0.5 |
| **58** | 9.75 | 7.5 | 5.25 | 4 | 3.5 | 3 | 2 | 1.5 | 1 | 0.25 |
| **59** | 9.75 | 7.5 | 5.25 | 4 | 3.5 | 3 | 2 | 1.5 | 1 | 0.25 |
| **60** | 9.75 | 7.5 | 5 | 4 | 3.5 | 3 | 2 | 1.5 | 1 | 0.25 |
| **61** | 9.5 | 7.5 | 4.75 | 3.75 | 3.5 | 3 | 2 | 1.25 | 1 | 0.25 |
| **62** | 9.5 | 7.5 | 4.75 | 3.75 | 3.5 | 3 | 2 | 1.25 | 0.75 | 0.25 |
| **63** | 9.5 | 7.25 | 4.75 | 3.5 | 3.25 | 3 | 2 | 1.25 | 0.75 | 0.25 |

| | | | | | | | | | | |
|---|---|---|---|---|---|---|---|---|---|---|
| **64** | 9.25 | 7 | 4.75 | 3.25 | 3.25 | 3 | 2 | 1.25 | 0.75 | 0.25 |
| **65** | 9.25 | 7 | 4.5 | 3.25 | 3.25 | 2.75 | 2 | 1.25 | 0.75 | 0.25 |
| **66** | 9 | 7 | 4.5 | 3 | 3.25 | 2.75 | 1.75 | 1.25 | 0.75 | 0.25 |
| **67** | 9 | 6.75 | 4.5 | 3 | 3.25 | 2.75 | 1.75 | 1.25 | 0.75 | 0.25 |
| **68** | 8.75 | 6.5 | 4.5 | 2.75 | 3.25 | 2.75 | 1.75 | 1.25 | 0.75 | 0.25 |
| **69** | 8.5 | 6.5 | 4.5 | 2.5 | 3 | 2.5 | 1.75 | 1.25 | 0.75 | 0.25 |
| **70** | 8.5 | 6.5 | 4.5 | 2.5 | 3 | 2.5 | 1.75 | 1 | 0.5 | 0.25 |

**ESI. Table 5:** Mean and Standard Deviation (SD) of 7 healthy samples in ESI. Table 5 above, diluted to different concentration between 5 to 50 % Hct The tabulated data is the 99% confidence interval of every concentration (7×10=70 test results each).

| **SR. No.** | **1** | **2** | **3** | **4** | **5** | **6** | **7** | **8** | **9** | **10** |
|---|---|---|---|---|---|---|---|---|---|---|
| **Conc. (%)** | 5 | 10 | 15 | 20 | 25 | 30 | 35 | 40 | 45 | 50 |
| **Average l(t=5min), (mm)** | 11.25 | 9.12 | 7.00 | 5.13 | 5.19 | 3.45 | 2.64 | 2.11 | 1.49 | 0.84 |
| **Standard Deviation (SD)** | 2.07 | 1.58 | 2.27 | 1.49 | 2.26 | 0.61 | 0.69 | 0.82 | 0.52 | 0.51 |
| **Lower Limit** | 10.62 | 8.64 | 6.30 | 4.67 | 4.50 | 3.27 | 2.43 | 1.86 | 1.32 | 0.68 |
| **Upper Limit** | 11.89 | 9.61 | 7.70 | 5.58 | 5.89 | 3.64 | 2.85 | 2.37 | 1.65 | 0.99 |

**ESI. Table 6:** Data set of 71 blood samples used to obtain the length traversed in the paper device without any dilution or sample treatment. Samples with % Hct values greater than 33% were mostly obtained from B.C Roy Technology Hospital, Indian Institute of Technology, Kharagpur, and the samples below 33% Hct were obtained from Institute of Haematology and Transfusion Medicine, Calcutta Medical College, Kolkata.

| **SR. No.** | **% Hct** | **Length (mm)** | **Obtained Interval (%)** |
|---|---|---|---|
| **1** | 11.3 | 9.25 | **10-20** |
| **2** | 13.8 | 11.5 | **10-20** |
| **3** | 15.2 | 10.75 | **10-20** |
| **4** | 16.3 | 16.25 | **10-20** |
| **5** | 16.7 | 7.5 | **10-20** |
| **6** | 17.6 | 8 | **10-20** |
| **7** | 19.2 | 6.75 | **10-20** |
| **8** | 19.5 | 8 | **10-20** |
| **9** | 19.5 | 12 | **10-20** |
| **10** | 21.1 | 5.5 | **20-30** |
| **11** | 21.3 | 6 | **20-30** |
| **12** | 23.1 | 4 | **20-30** |
| **13** | 23.5 | 5.5 | **20-30** |

| **SR. No.** | **% Hct** | **Length (mm)** | **Obtained Interval (%)** |
|---|---|---|---|
| **37** | 34.6 | 3.75 | **30-40** |
| **38** | 34.7 | 2.75 | **30-40** |
| **39** | 34.8 | 2.25 | **30-40** |
| **40** | 35 | 2.5 | **30-40** |
| **41** | 35.3 | 1.75 | **30-40** |
| **42** | 35.7 | 3.25 | **30-40** |
| **43** | 35.8 | 3.25 | **30-40** |
| **44** | 36 | 2.25 | **30-40** |
| **45** | 36.6 | 3.25 | **30-40** |
| **46** | 36.7 | 2.5 | **30-40** |
| **47** | 36.7 | 2.75 | **30-40** |
| **48** | 36.9 | 2.5 | **30-40** |
| **49** | 37.5 | 1.5 | **30-40** |

| | | | |
|---|---|---|---|
| **14** | 24 | 4 | **20-30** |
| **15** | 24.3 | 5.75 | **20-30** |
| **16** | 25.4 | 8.25 | **20-30** |
| **17** | 26.5 | 4.25 | **20-30** |
| **18** | 26.6 | 10 | **20-30** |
| **19** | 26.9 | 3.75 | **20-30** |
| **20** | 28 | 4 | **20-30** |
| **21** | 28.3 | 5 | **20-30** |
| **22** | 28.5 | 4.25 | **20-30** |
| **23** | 28.6 | 3.5 | **20-30** |
| **24** | 28.6 | 6.5 | **20-30** |
| **25** | 28.7 | 4.25 | **20-30** |
| **26** | 28.8 | 3.5 | **20-30** |
| **27** | 29.1 | 4 | **20-30** |
| **28** | 29.4 | 5.5 | **20-30** |
| **29** | 29.6 | 3.75 | **20-30** |
| **30** | 30.1 | 2.75 | **30-40** |
| **31** | 30.3 | 5.5 | **30-40** |
| **32** | 30.9 | 2.75 | **30-40** |
| **33** | 31.9 | 3.5 | **30-40** |
| **34** | 31.9 | 3.5 | **30-40** |
| **35** | 33 | 2.5 | **30-40** |
| **36** | 34.2 | 2.25 | **30-40** |

| | | | |
|---|---|---|---|
| **50** | 38.7 | 2.25 | **30-40** |
| **51** | 38.9 | 1.5 | **30-40** |
| **52** | 39.6 | 1.75 | **30-40** |
| **53** | 40.8 | 1.5 | **40-50** |
| **54** | 41.5 | 2 | **40-50** |
| **55** | 41.8 | 1 | **40-50** |
| **56** | 42.4 | 2.25 | **40-50** |
| **57** | 42.4 | 1.25 | **40-50** |
| **58** | 42.6 | 0.75 | **40-50** |
| **59** | 43 | 2 | **40-50** |
| **60** | 44.5 | 2.25 | **40-50** |
| **61** | 44.9 | 1.5 | **40-50** |
| **62** | 45 | 0.5 | **40-50** |
| **63** | 45.1 | 1.5 | **40-50** |
| **64** | 45.4 | 0.5 | **40-50** |
| **65** | 45.7 | 0.25 | **40-50** |
| **66** | 46.9 | 0.5 | **40-50** |
| **67** | 47.2 | 1.5 | **40-50** |
| **68** | 47.4 | 1.25 | **40-50** |
| **69** | 48.3 | 0.25 | **40-50** |
| **70** | 49.9 | 0.75 | **40-50** |
| **71** | 50.3 | 0 | **40-50** |

**ESI. Table 7:** Results of blind test in the fast-flowing paper device. Data represents the length(mm) traversed for every randomly selected blood sample's %Hct and the obtained interval for the corresponding length. All the samples were acquired from Institute of Haematology and Transfusion Medicine, Calcutta Medical College, Kolkata.

| **SR. NO.** | **WBC** | **RBC** | **HGB** | **HCT** | **PLT** | **LENGTH (mm)** | **Obtained Interval (%)** |
|---|---|---|---|---|---|---|---|
| **1** | 69.32+ | 2.52 | 8.1 | 36.5 | 118 | 1.75 | 40-50 |
| **2** | 18.67+ | 3.63 | 11.1 | 34 | 80 | 3.25 | 30-40 |
| **3** | 2.3- | 1.79- | 5.4- | 15.0 | 8- | 8.25 | 10-20 |
| **4** | 23.37+ | 4.41 | 9.8 | 32.4 | 230 | 3.5 | 30-40 |
| **5** | 2.90- | 1.50- | 4.6- | 13.7- | 4- | 5.75 | 10-20 |
| **6** | 7.83 | 5.44 | 12.1 | 40.1 | 29- | 3.75 | 40-50 |
| **7** | 5.99 | 3.93 | 13.6 | 40.6 | 121 | 2.5 | 40-50 |
| **8** | 6.85 | 1.47 | 2.3- | 10 | 149 | 12 | 10-20 |
| **9** | 13.11 | 4.77 | 13.2 | 42.4 | 644+ | 2 | 40-50 |
| **10** | 3.08 | 4.37 | 13.4 | 39.9 | 151 | 3 | 30-40 |
| **11** | 46.3+ | 3.10 | 11.1 | 23.8 | 90 | 5 | 20-30 |
| **12** | 9.05 | 3.54 | 12.4 | 37.1 | 148 | 3 | 30-40 |

| | | | | | | | |
|---|---|---|---|---|---|---|---|
| **13** | 23.98+ | 4.93 | 12.4 | 41.3 | 941+ | 3.25 | 40-50 |
| **14** | 2.27- | 2.35- | 6.9- | 20.9- | 9- | 9.5 | 20-30 |
| **15** | 5.99 | 3.93 | 13.3 | 28.7 | 86 | 3.25 | 20-30 |
| **16** | 5.67 | 4.5 | 10.9 | 38.6 | 183 | 3.5 | 30-40 |
| **17** | 14.62 | 2.83 | 9.7 | 30.7 | 398 | 4 | 30-40 |
| **18** | 10.35 | 6.46+ | 15 | 48.1 | 253 | 1.5 | 40-50 |
| **19** | 6.73 | 4.61 | 12.8 | 39.1 | 171 | 2.5 | 40-50 |
| **20** | 8.57 | 3.22 | 6.6- | 21.4 | 215 | 8.5 | 20-30 |
| **21** | 16.99+ | 3.68 | 8.4 | 25.2- | 265 | 4 | 20-30 |
| **22** | 26.8+ | 4.3 | 12.7 | 42.3 | 347 | 3.25 | 30-40 |
| **23** | 6.52 | 4.87 | 13.9 | 43.7 | 135 | 2.5 | 40-50 |
| **24** | 3.53 | 3.48 | 9.7 | 31.1 | 74 | 4.25 | 30-40 |
| **25** | 6.00 | 5.73+ | 14.1 | 43.2 | 148 | 2.25 | 40-50 |
| **26** | 2.9- | 1.5- | 4.6- | 13.7- | 4- | 7.75 | 10-20 |
| **27** | 5.87 | 3.57 | 10.3 | 32.2 | 86 | 2.5 | 30-40 |
| **28** | 4.04 | 2.06- | 7.9- | 25.4- | 106 | 3.25 | 30-40 |
| **29** | 22.13+ | 3.71 | 9.9 | 31.4 | 236 | 2.75 | 30-40 |
| **30** | 13.25 | 4.23 | 10.5 | 32.8 | 259 | 2.75 | 30-40 |
| **31** | 247.05+ | 2.79 | 4.6- | 16.3 | 97 | 5 | 20-30 |
| **32** | 2.3- | 2.2 | 7.6- | 22.2- | 39- | 6.75 | 20-30 |
| **33** | 3.39 | 3.39 | 9.3 | 26.6 | 5- | 4.25 | 20-30 |
| **34** | 5.76 | 2.76 | 8.4 | 27.1 | 114 | 4 | 20-30 |
| **35** | 45.82+ | 3.89 | 12.8 | 37.1 | 114 | 2.75 | 30-40 |
| **36** | 8.02 | 3.06 | 7.3- | 25.3- | 412+ | 3 | 30-40 |
| **37** | 8.83 | 4.54 | 7.4- | 26.2 | 293 | 4 | 20-30 |
| **38** | 12.54 | 4.63 | 12.6 | 39.9 | 135 | 3.5 | 30-40 |
| **39** | 8.37 | 4.30 | 11.4 | 35.7 | 260 | 2.5 | 30-40 |
| **40** | 6.21 | 3.41 | 10.8 | 34.3 | 150 | 2.5 | 30-40 |
| **41** | 8.88 | 4.95 | 15.7 | 48.1 | 143 | 1.5 | 40-50 |
| **42** | 11.97 | 3.09 | 6.3- | 21.5 | 402+ | 4.75 | 20-30 |
| **43** | 4.84 | 2.69 | 8.8 | 31.7 | 93 | 3.25 | 30-40 |
| **44** | 6.38 | 4.69 | 12.6 | 39.5 | 149 | 3 | 30-40 |
| **45** | 9.2 | 3.64 | 9.6 | 29.7 | 157 | 4.75 | 20-30 |
| **46** | 6.52 | 3.28 | 9.9 | 33.6 | 42- | 3.5 | 30-40 |
| **47** | 3.4 | 2.00- | 5.9- | 17.9 | 3- | 6.5 | 20-30 |
| **48** | 3.6 | 2.06- | 4.8- | 19.8 | 3- | 6.75 | 20-30 |
| **49** | 8.13 | 4.57 | 10.9 | 36.4 | 122 | 3.25 | 30-40 |
| **50** | 6.8 | 3.4 | 11.1 | 33 | 43 | 2.75 | 30-40 |
| **51** | 5.65 | 3.27 | 12.2 | 36.6 | 112 | 2.75 | 30-40 |
| **52** | 6.24 | 4.58 | 12.4 | 39.6 | 143 | 2.5 | 30-40 |
| **53** | 12.91 | 3.68 | 10.4 | 30.7 | 162 | 3.25 | 30-40 |
| **54** | 7.06 | 3.82 | 7.8- | 24.7 | 145 | 3.5 | 20-30 |

** The images of highlighted Hct values in yellow colour are given in table below.

The blind Hct detection using our fold induced paper channels showed an estimation accuracy of 90.74%, highlighting the clinical relevance and reliability of our paper folded channels for designing biomedical devices.

**ESI. Table 8:** Images representing randomly selected and performed blind test on two more simultaneous devices for whole blood samples with %Hct chosen from each interval.

| % HEMATOCRIT | | | |
|---|---|---|---|
| **10-20 %** | **20-30 %** | **30-40 %** | **40-50 %** |
| 15 | 23.8 | 30.7 | 48.1 |
| 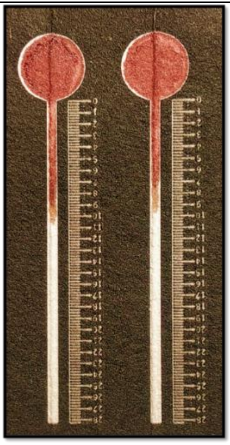 | 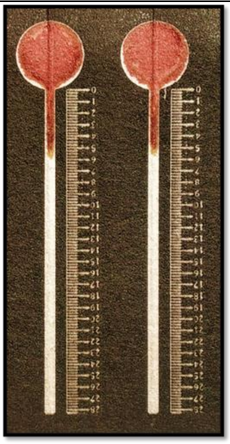 | 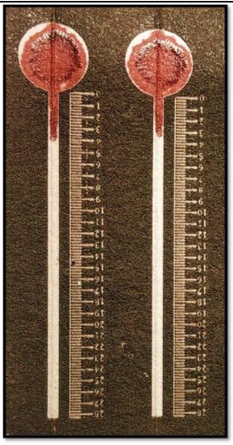 | 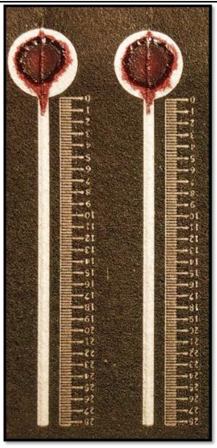 |
